# Incomplete developmental silencing of cancer-testis antigen BORIS in humanized mouse model promotes cancer susceptibility

**DOI:** 10.64898/2025.12.24.696438

**Authors:** Emma Price, Elena M. Pugacheva, Dharmendra N. Bhatt, Yon J. Ji, Sheila Yeboah, Arielle Scott, Dmitri Loukinov, Victor Lobanenkov

## Abstract

Cancer-testis antigens are genes normally restricted to the germline but aberrantly activated in many cancers, where their role in tumorigenesis remains unclear. BORIS (CTCFL), a testis-specific paralog of the chromatin organizer CTCF, is one such antigen that shows limited conservation between humans and mice, complicating *in vivo* functional studies. Here, we generated humanized mice in which both alleles of the endogenous *Boris* locus are replaced with the full-length human *BORIS* gene, including its highly diverged *cis*-regulatory elements. These fully humanized mice maintain normal fertility, indicating accurate germline expression and preserved BORIS function despite evolutionary divergence. However, unlike the strictly testis-specific expression of mouse *Boris*, human *BORIS* escapes complete somatic silencing, producing mosaic expression in a minority of mouse somatic cells. This ectopic expression is associated with reduced survival, increased tumor incidence and a shift of tumor spectrum toward aggressive lymphomas. Transcriptomic and chromatin profiling revealed that ectopic human BORIS reactivates testes-specific genes in the soma, including regulators of meiosis and DNA repair, through direct chromatin binding. This transcriptional reprogramming was consistent across tissues and clonal cell lines, revealing a dominant tissue-independent gene activation program. These findings demonstrate that when human BORIS escapes epigenetic silencing in somatic cells, it connects aberrant germline gene activation with increased cancer susceptibility *in vivo*.

## Introduction

BORIS (CTCFL, Brother of Regulator of Imprinted Sites) is a testis-specific paralog of the ubiquitous chromatin organizer CTCF, sharing its highly conserved 11-zinc-finger DNA-binding domain, but diverging substantially in N- and C-termini^1,2^. While CTCF is essential for 3D genome organization and expressed in all cell types^3^, mammalian BORIS exhibits restricted expression exclusively in male germ cells during spermatogenesis^2,4^, where both CTCF and BORIS co-localize at genomic sites and cooperatively regulate germline-specific transcriptional programs^5^.

Germline-restricted transcription is frequently reactivated in human cancers, where BORIS is also often aberrantly expressed, leading to its classification as a cancer-testis antigen (CTA)^1,6,7^. CTAs represent a unique class of tumor-associated antigens that are normally restricted to immune-privileged germ cells but can also be ectopically expressed in tumors, making them attractive targets for cancer immunotherapy^8–10^. Several CTAs, including NY-ESO-1 and MAGE-A3, have advanced to clinical trials with promising results^11,12^, yet the specific mechanisms of how CTAs become activated and if their activation contributes to tumorigenesis remain poorly understood outside of *in vitro* models. Ectopic BORIS expression in cell lines results in epigenetic reprogramming of CTCF binding sites^13^ and activation of oncogenic transcriptional programs^14–18^, but whether BORIS actively drives transformation or simply represents a downstream consequence of cellular perturbations has not yet been tested in physiologically relevant models. It is important to note that cancer cell lines, despite their widespread use, inherently carry abnormal karyotypes, accumulated mutations and genomic instability that may influence experimental outcomes ^19^.

BORIS arose through a whole gene duplication of CTCF, creating a paralog that has since undergone rapid evolutionary divergence in vertebrates^2,20^. BORIS exhibits major changes in its N- and C-terminal domains while retaining a highly conserved 11-zinc-finger DNA-binding domain, nearly identical to that of CTCF. This conservation enables both proteins to bind the same CTCF target sites (CTSs)^2^, raising the possibility of competitive binding. However, *in vivo* studies suggest a more complex interaction. Instead of direct competition, CTCF and BORIS often co-bind at specialized clustered DNA elements called 2xCTSes, which contain two or more adjacent CTCF motifs^5,7^. Notably, BORIS does not bind most single CTCF motifs^21^, likely due to its limited ability to recruit chromatin remodelers like SMARCA5, which CTCF uses to reposition nucleosomes and facilitate DNA binding^22^. Across vertebrate evolution, BORIS expression became restricted to the male germline, coinciding with the evolution of genomic imprinting in mammals^20^, a process primarily regulated through CTCF binding sites ^23^. However, the extent of BORIS requirement for fertility differs strikingly between species, with primate BORIS showing evidence of expanded function compared to rodent orthologs^24,25^.

While BORIS knockout (KO) mice exhibit only subfertility with partial spermatogenic defects^4,26,27^, a homozygous loss-of-function mutation in human *BORIS* causes complete sterility^28^, indicating a more critical role in human reproduction. Human *BORIS* utilizes three alternative promoters generating 22 transcript isoforms, including expression from a CpG island-containing promoter (named promoter B), whereas mouse *Boris* produces only two isoforms from a single promoter^29,30^, highlighting a substantial difference in regulatory complexity between species. In macaque testes, BORIS exhibits higher expression levels and occupies significantly more genomic sites than in mouse, including co-localization with meiotic cohesin subunits^25^, suggesting expanded regulatory capacity in primates. These species-specific differences in BORIS expression, genomic occupancy, and fertility requirements suggest that human BORIS has evolved functions that cannot be fully recapitulated in standard mouse models, limiting our ability to study its function in fertility and disease.

Despite growing interest in human BORIS as both a cancer biomarker and therapeutic target, fundamental questions about its pathological function remain unanswered. Functional studies have been constrained by limitations of existing models. While mouse knockout studies reveal germline requirements they cannot address human-specific BORIS function, while transgenic human BORIS overexpression in cancer cell lines produces artificially high levels that may not reflect endogenous regulation or mosaic expression patterns often observed in human tumors^31,32^. Moreover, the substantial sequence divergence between human BORIS (hBORIS) and mouse BORIS (mBORIS) proteins—particularly in N- and C-terminal regions – severely limits the translatability of functional insights gained from mice studies to human biology.

Cell line studies have demonstrated that ectopic BORIS expression can reprogram CTCF-occupied sites and activate gene expression programs ^13,14^, but whether this ectopic human BORIS expression actively drives tumorigenesis *in vivo* or simply represents a downstream consequence of transformation has not been directly tested. Furthermore, mechanisms by which CTAs like BORIS contribute to cellular transformation and tumor progression *in vivo* remain poorly defined, hampering efforts to target these proteins therapeutically. Addressing specific human CTA-related questions relating to functional and regulatory divergence in both reproductive and pathological contexts is a complex task and requires a physiologically relevant mouse model that mimics endogenous regulatory control in both germline and pathogenic contexts^33^. To address this directly, we created a unique humanized BORIS mouse with preserved human *cis*-regulatory elements.

## Results

### Generation and Validation of a Humanized *BORIS* Mouse Model

To develop a physiologically relevant model to study hBORIS, we replaced the endogenous mouse *Boris* locus with the complete human *BORIS* locus, including upstream and downstream *cis*-regulatory elements (Fig 1A). This approach was motivated by substantial structural and regulatory divergence between species (Fig S1A-C). While mouse *Boris* is transcribed from a single promoter producing two isoforms (Fig S1C), human *BORIS* utilizes three alternative promoters (named promoter A, B, and C) (Fig S1B) generating 22 transcripts encoding 16 protein isoforms^30^ (Fig S1A). Canonical human BORIS in human testes is expressed from promoter B (Fig S1B), which contains a CpG island and flanking H3K4me1-marked enhancers (Fig S1A-C). We retained these human-specific regulatory elements to ensure proper *in vivo* expression (Fig 1A, Fig S1A).

**Figure 1.**
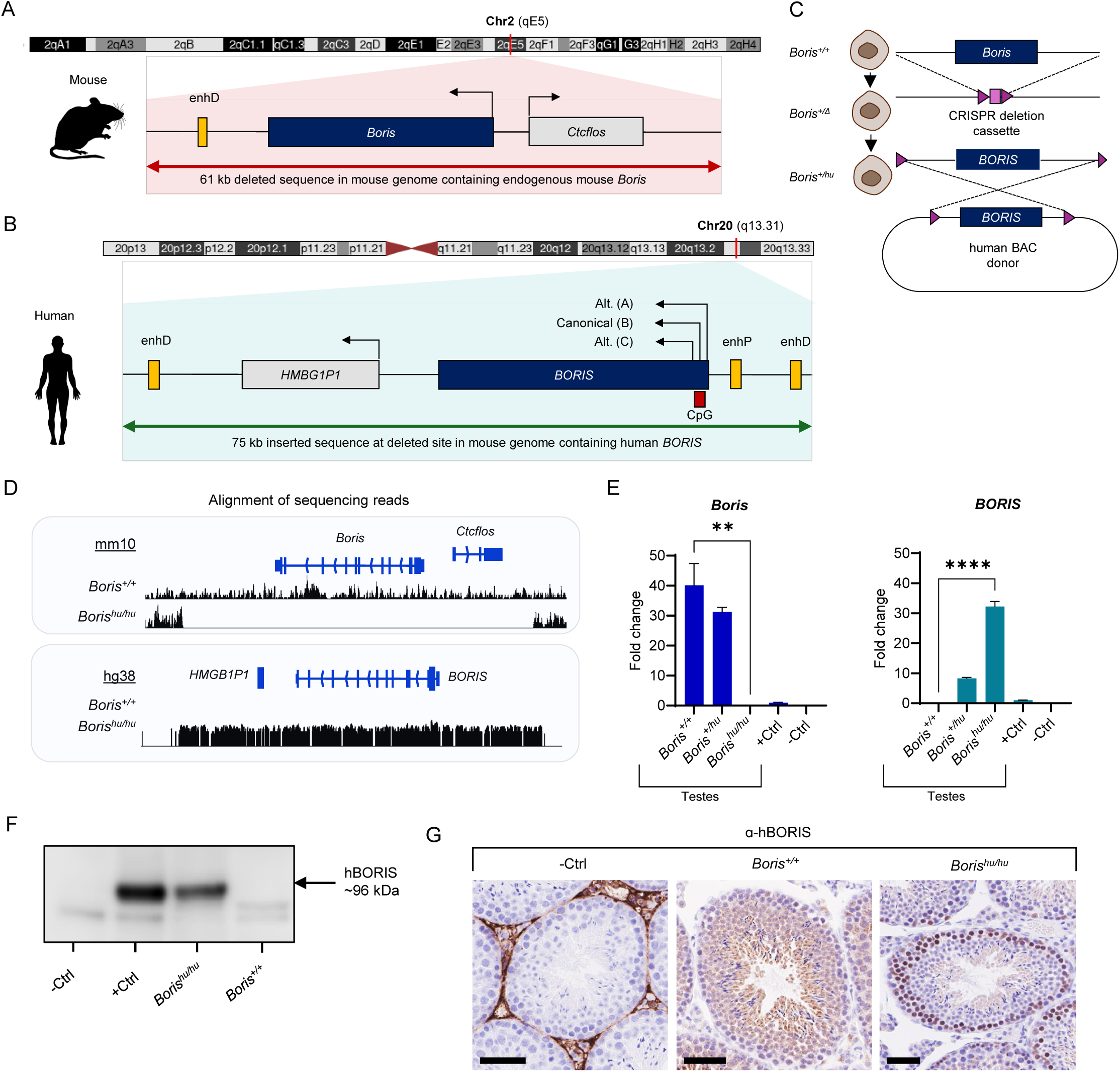
CRISPR–RMGR–mediated humanization of the endogenous mouse *Boris* locus and validation in mouse testes. (A) Schematic of the endogenous mouse *Boris* locus on chromosome 2 (qE5). A 61-kb interval (chr2:173,077,709–173,138,357, mm10) was deleted to remove the entire mouse gene and its regulatory region. The distal enhancer (enhD) is shown in yellow. See Fig. S1 for extended annotation. (B) Schematic of the human *BORIS* locus on chromosome 20 (q13.31). A 75-kb genomic fragment (chr20:57,471,893–57,547,144, hg38), containing promoters A, B, and C, proximal and distal enhancers (enhP, enhD) (yellow), and the CpG island at promoter B (red), was inserted into the deleted mouse locus. See Fig. S1 for extended annotation. (C) Overview of the humanization strategy. CRISPR/Cas9 was used to excise the 61-kb mouse *Boris* sequence (panel A), followed by recombinase-mediated genomic replacement (RMGR) to insert the full 75-kb human *BORIS* sequence (panel B) from a BAC donor construct. Additional construct details are shown in Fig. S2A–B. (D) Genome browser validation of allele replacement. Upper: sequencing reads from *Boris^+/+^* testes align uniquely to the mouse mm10 locus, whereas reads from *Boris^hu/hu^* testes show complete absence of alignment across the deleted interval. Lower: reads from *Boris^hu/hu^* testes align across the inserted human locus (hg38), while *Boris^+/+^* reads do not, confirming precise deletion of the mouse sequence and insertion of the human locus. (E) qRT–PCR quantification of mouse *Boris* and human *BORIS* mRNA in testes from *Boris^+/+^, Boris^+/hu^,* and *Boris^hu/hu^* mice (n = 3 biological replicates per genotype). Statistical testing by t-test; p < 0.05. +Ctrl: *Boris^+/hu^* mESCs (panel C), which express both human and mouse transcripts (see Fig. 2). –Ctrl: *Boris^−/−^* knockout mouse testes, which do not express either *Boris* or *BORIS*. (F) Representative Western blot using a hBORIS–specific antibody shows robust protein expression in *Boris^hu/hu^* testes but no detectable signal in *Boris^+/+^* testes. +Ctrl: OVCAR8 cells, which express high levels of human BORIS. –Ctrl: OVCAR3 cells, which do not express human BORIS. (G) Immunohistochemistry demonstrates hBORIS protein nuclear staining in spermatogonia and spermatocytes of *Boris^hu/hu^* testes. No signal is detected in *Boris^+/+^* testes, confirming antibody specificity. Scale bar, 50 μm. –Ctrl: *Boris^hu/hu^* testes processed without secondary antibody.

In mouse embryonic stem cells (mESCs), we removed a 61 kb region encompassing the endogenous mouse *Boris* locus on chromosome 2 (chr2:173,077,709–173,138,357) using CRISPR, then inserted a 75 kb human sequence containing the complete *BORIS* gene from chromosome 20 (chr20:57,471,893–57,547,144) via recombinase-mediated genomic replacement (RMGR) (Fig 1A-C, Fig S2A-B). Long-read PacBio sequencing confirmed transgene sequence integrity (Table S1). Heterozygous (*Boris^+/hu^*) mESCs were used to establish founder mice, which were bred to generate homozygous humanized (*Boris^hu/hu^*) and wildtype (*Boris^+/+^*) littermates (Fig 1C, Fig S2C). Genotyping was confirmed by PCR (Fig S2D) with offspring born at expected Mendelian ratios (Fig S2E). Alignment of sequencing reads validated locus replacement in *Boris^hu/hu^* testes, with *BORIS*-aligned reads mapping exclusively to the human locus (chr20), and zero reads mapping at the deleted mouse locus (chr2), whereas *Boris^+/+^* controls showed mouse-specific *Boris*-aligned mapping (Fig 1D).

To directly compare expression of human *BORIS* and mouse *Boris* orthologs, we performed qRT-PCR using heterozygous *Boris^+/hu^* mESCs as a reference sample for both genes (Fig 1E). In *Boris^hu/hu^* testes, human *BORIS* mRNA was expressed at 32-fold relative to the *Boris^+/hu^* mESC reference, comparable to the 40-fold expression of mouse *Boris* in *Boris^+/+^* testes (Fig 1E). Mouse *Boris* was not detectable in *Boris^hu/hu^* testes, and no human *BORIS* was detectable in *Boris^+/+^* testes, validating locus-specific expression and primer specificity. These data demonstrate that the humanized *BORIS* allele produces mRNA at physiologically relevant levels comparable to the endogenous mouse *Boris* ortholog.

Western blot confirmed human BORIS protein expression in *Boris^hu/hu^* testes (Fig 1F). Immunohistochemistry (IHC) further revealed hBORIS nuclear localization in *Bors^hu/hu^* spermatogonia and primary spermatocytes, with appropriate absence in later-stage germ cells, consistent with the expected expression window during spermatogenesis^5^ (Fig 1G). RNA-seq confirmed that *Boris^hu/hu^* testes express the full-length canonical human BORIS transcript from promoter B, mirroring known expression in human testes (Fig S2F). Together, these data demonstrate that the human *cis*-regulatory elements included in the humanized locus retain their function in the *Boris^hu/hu^* mouse genomic landscape, supporting appropriate cell type-specific hBORIS expression and developmental timing in the murine germline.

### Human BORIS Expression Detected in Somatic Tissues

Ectopic hBORIS activation outside of the germline has been reported in multiple human cancers^31,34^, raising questions about whether the humanized *BORIS* locus would remain repressed/silent in somatic tissues within the humanized model. We examined human *BORIS* expression across multiple tissues in *Boris^hu/hu^* mice. As expected, mouse *Boris* remained completely undetectable in all somatic tissues in *Boris^+/+^* mice (Fig 2A). Unexpectedly, human *BORIS* mRNA was detected in all somatic tissues in *Boris^hu/hu^* mice; at robust but variable levels (Fig 2A). RNA-seq confirmed that several tissues expressed the full-length canonical human *BORIS* transcript (Fig S3A). Promoter B (Fig S1B), drove the majority of transcription across somatic tissues, though promoter C showed activity in liver, and alternative splicing variants were detected in several tissues (Fig S3A). This alternate promoter usage pattern is consistent with reports from human cancers^29^. Promoter A remained inactive in all tissues examined (Fig S3A).

**Figure 2.**
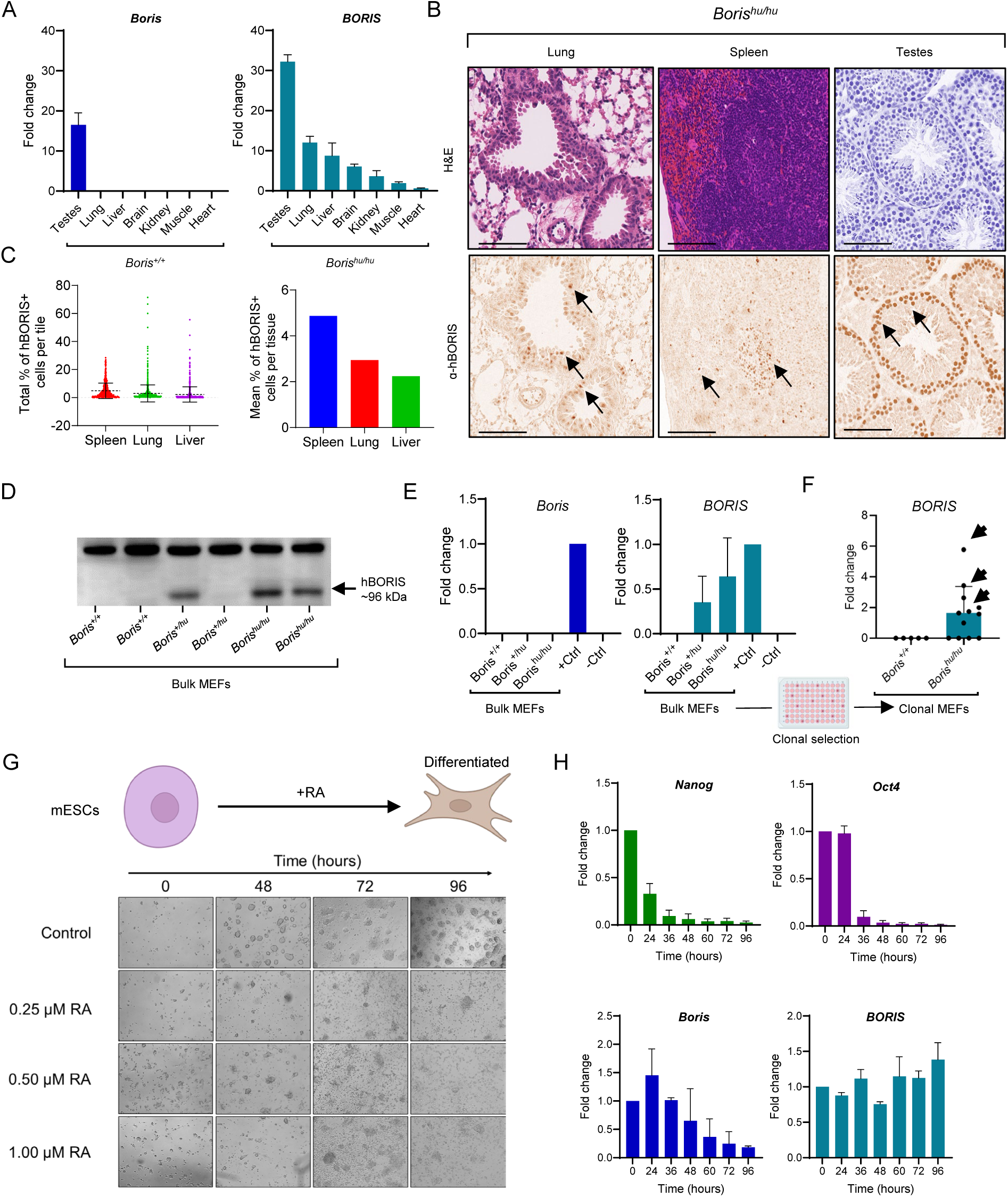
Human *BORIS* escapes epigenetic repression across diverse somatic contexts. (A) qRT–PCR quantification of endogenous mouse *Boris* in *Boris^+/+^* tissues and human *BORIS* mRNA in *Boris^hu/hu^* somatic tissues (n = 3 biological replicates; 3 technical replicates; age 16 weeks). Species-specific primers were used. Data are normalized to *Gapdh* and presented as ΔΔCt values relative to testes samples. (B) Immunohistochemistry using a human BORIS–Ab reveals rare but reproducible hBORIS-positive nuclei in *Boris^hu/hu^* somatic tissues. Positive cells (brown DAB signal, arrows) were identified in spleen, liver, and brain. Counterstaining with hematoxylin confirms intact tissue architecture; matched H&E sections are shown for comparison. Scale bars, 50 μm. (C) Quantification of hBORIS-positive cells across somatic tissues. Left: automated nuclear segmentation and DAB-positive cell classification, reporting the percentage of hBORIS-positive cells per tile (see Fig. S3). Error bars; ± SEM. Right: whole-section quantification showing mean hBORIS-positive frequency per tissue (± SEM). (D) Western blot analysis of bulk MEF lysates derived from *Boris^+/+^*, *Boris^+/hu^*, and *Boris^hu/hu^* embryos. A human BORIS–specific antibody detects the expected ~96-kDa hBORIS band exclusively in *Boris^+/hu^* and *Boris^hu/hu^* MEFs, with no detectable signal in *Boris^+/+^* cultures. The upper band present in all samples represents a non-specific antibody reactivity and serves as an internal reference for protein transfer. (E) qRT–PCR quantification of endogenous mouse *Boris* in *Boris^+/+^* bulk MEFs and human *BORIS* mRNA in *Boris^hu/hu^* bulk MEFs (n = 2 biological replicates, 3 technical replicates). Species-specific primers were used. Data are normalized to *Gapdh* and presented as ΔΔCt values. (F) qRT–PCR quantification of human *BORIS* mRNA in clonally expanded *Boris^hu/hu^* MEFs (n = 12 biological replicates, 3 technical replicates). Species-specific primers were used. Data are normalized to *Gapdh* and presented as ΔΔCt value. (G) Schematic of retinoic acid (RA)–induced differentiation protocol used for heterozygous *Boris^+/hu^* mESCs. Cells were treated with 1 μM RA for 96 h, with samples collected at defined time points. Representative bright-field images illustrate the transition from compact pluripotent colonies to flattened, differentiated monolayers. (H) qRT–PCR analysis of RA-differentiated *Boris^+/hu^* mESCs demonstrating that endogenous mouse Boris is downregulated in coordination with pluripotency markers (*Nanog, Oct4*) during differentiation, whereas human *BORIS* fails to undergo transcriptional silencing. Data are normalized to *Gapdh* and plotted ΔΔCt values relative to day-0 undifferentiated cells (n = 2 biological replicates 3 technical replicates).

IHC across several tissues revealed that hBORIS was not uniformly expressed in all cells but instead showed a striking mosaic pattern (Fig 2B, Fig S3B). To quantify this heterogeneity, tissue sections were divided into equal-sized tiles and scored for hBORIS-positive cells (Fig S3C). This analysis showed dense hBORIS staining indicative of localized hBORIS-positive cell clusters, but most areas contained very few or zero positive cells indicative of very sporadic activation (Fig 2C, Fig S3C). Quantification across tissues showed on average 2-5% of cells scored positive for hBORIS expression at the protein level (Fig 2C), with variation between tissue types. Within this mosaic pattern, anatomical distribution varied by organ (Fig S3B): in spleen, BORIS-positive cells appeared in both red and white pulp with hBORIS enrichment in germinal centers; in lung, positive cells were observed in bronchiolar epithelium, alveolar pneumocytes, and lymphocytic infiltrates; and in liver, scattered staining appeared particularly in perivascular zones.

### Human BORIS Silencing is Incomplete During Lineage Commitment

To examine regulation of the humanized *BORIS* locus in a more homogeneous system, we generated mouse embryonic fibroblast (MEF) cell lines from *Boris^hu/hu^* and *Boris^+/+^* E13.5 embryos. Consistent with previous reports, endogenous mouse *Boris* was not detected in wildtype MEFs, but human *BORIS* was robustly expressed in humanized MEFs (Fig 2D-E). We hypothesized that the failure to repress *BORIS* may occur during early embryonic development and persist throughout lifespan contributing to the mosaicism observed in IHC, as opposed to just repeated instances of ectopic activation in adult somatic tissues (Fig 2E). Next, we isolated clonal MEF lines from parental cultures via limiting dilution to assess the heterogeneity of human *BORIS* expression between genetically identical cells (Fig 2D-E). Strikingly, human *BORIS* expression was indeed variable across genetically identical clones (Fig 2F). While some clones showed robust *BORIS* expression, others showed no detectable *BORIS* (Fig 2F). This highlights some degree of epigenetic variability in somatic cells whereby silencing is inefficient or unstable. The coexistence of hBORIS-positive and hBORIS-negative clones from the same *Boris^hu/hu^* genetic background demonstrates effectively that hBORIS *can* be silenced during development, but penetrance in this mouse strain is incomplete (Fig 2F). This cellular heterogeneity observed in culture, mirrors the mosaic expression patterns observed in *Boris^hu/hu^* somatic tissues (Fig 2B-C) and resembles the variable *BORIS* expression reported in human cancers, indicative of a mosaic epimutation in this mouse model^32^. For subsequent molecular profiling, we selected several hBORIS-expressing clones to support our dissection of hBORIS-driven transcriptional programs in the soma (Fig 2F).

BORIS is transiently expressed in stem cells and implicated in maintaining pluripotency^31,34^, therefore we leveraged heterozygous *Boris^+/hu^* mESCs generated during RMGR (Fig 1C) to profile both endogenous mouse *Boris* and human *BORIS* transcriptional regulation during retinoic acid (RA)-induced differentiation (Fig 2G). This served as a novel opportunity to directly test whether ectopic human *BORIS* expression may result from a failure to silence during early lineage commitment. As expected, pluripotency markers (*Oct4*, *Nanog*) and endogenous *Boris* were downregulated upon differentiation (Fig 2G-H), consistent with prior studies^35^. However, human *BORIS* remained detectable throughout the 96-hour differentiation time course, indicating incomplete silencing during exit from pluripotency (Fig 2G-H). Together, these findings suggest a degree of regulatory divergence between human *BORIS cis*-regulatory elements encoded within the humanized locus and somatic mouse trans-acting factors, enabling hBORIS to escape silencing in the soma. From our observations in both the germline and somatic tissues, two main questions were raised: (1) Does the substitution of mBORIS with hBORIS affect reproductive fitness in mice? (2) Does ectopic hBORIS activation have functional consequences in the soma?

### Human BORIS Supports Spermatogenesis in Mice

First, we assessed to what degree hBORIS retains functional conservation with mBORIS. Previous studies have established that mBORIS is important for fertility outcomes in male mice. Complete absence of mBORIS in *Boris^−/−^* knockout mice causes subfertility characterized by testicular atrophy, increased germ cell apoptosis, and reduced sperm counts evident at 90 days of age, ultimately resulting in reduced litter sizes^4,26^ (Fig 3D). This phenotype is dramatically exacerbated when combined with *Ctcf* haploinsufficiency, as double *Ctcf^+/−^ Boris^−/−^* mutants exhibit complete infertility due to severely disrupted transcriptional regulation during spermatogenesis^5^ (Fig 3D). These findings highlight the co-operative roles of mBORIS and CTCF in maintaining germline transcriptional programs. Notably, in humans, a loss-of-function mutation in hBORIS resulted in complete sterility rather than subfertility^28^, suggesting a greater evolutionary dependency on hBORIS for reproductive fitness. The substantial sequence divergence between hBORIS and mBORIS proteins (Fig S1D), combined with the regulatory differences observed in somatic contexts (Fig 2A-H), prompted us to test whether hBORIS can functionally replace mBORIS to support spermatogenesis.

**Figure 3.**
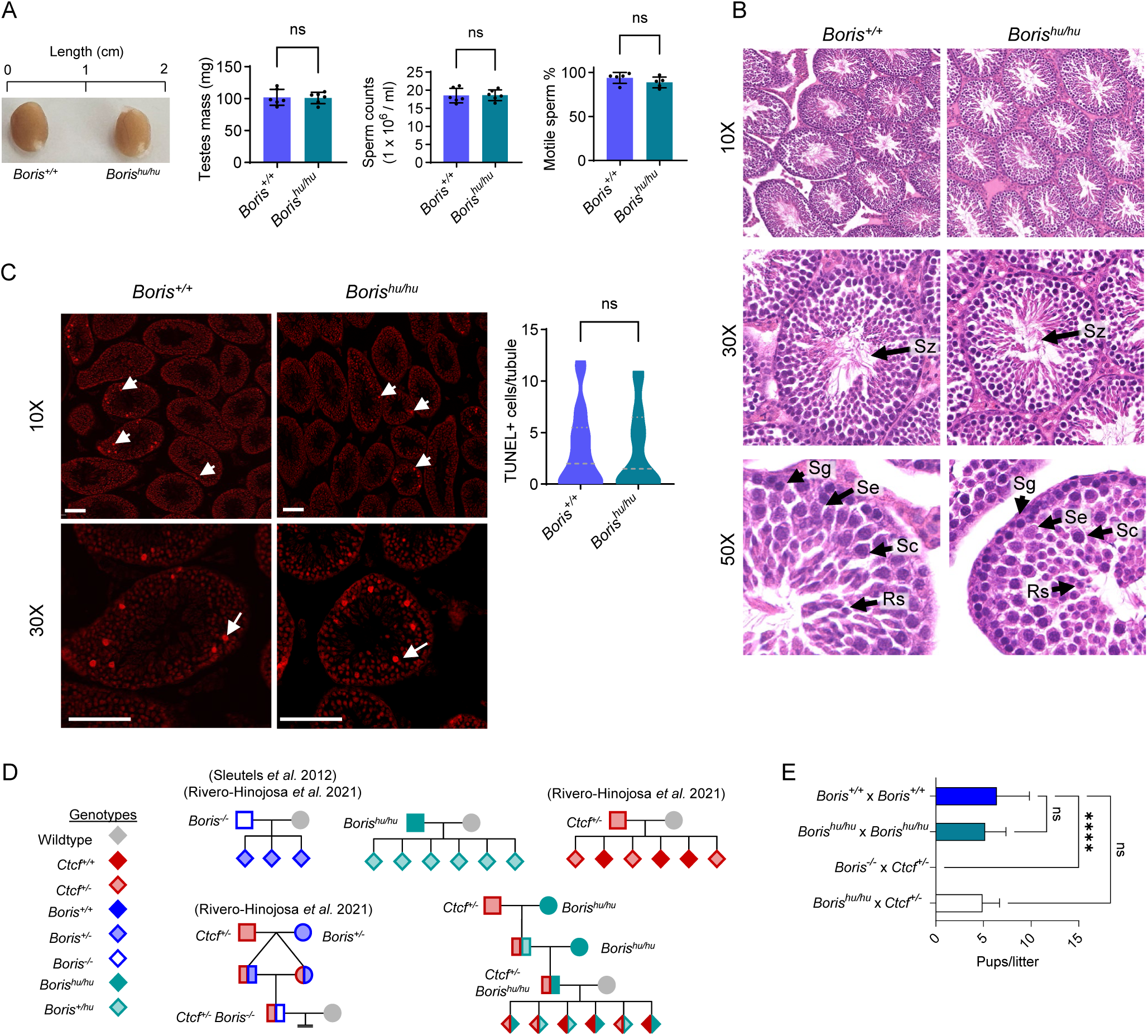
hBORIS in *Boris^hu/hu^* mice rescues subfertility phenotype associated with *Boris^−/−^* and Ctcf^+/−^ *Boris^−/−^* mutant mouse strains. (A) (Left to right) Testes size, testis mass, epididymal sperm counts, and sperm motility were quantified in Boris^+/+^ and Boris^hu/hu^ males aged 90 days (n = 5 biological replicates). Two-tailed unpaired t-tests; p < 0.05. (B) High-resolution H&E testes sections from Boris^+/+^ and Boris^hu/hu^ males aged 90 days show intact seminiferous tubule morphology with normal representation of spermatogonia (Sg), early spermatocytes (Se), spermatocytes (Sc), round spermatids (Rs), and spermatozoa (Sz). Published *Boris^−/−^* knockouts exhibit tubule atrophy and germ-cell loss at this age. Scale bar, 50 μm. (C) Representative TUNEL-stained sections (left) and quantification of TUNEL-positive cells per seminiferous tubule (right) in 90-day *Boris^+/+^* and *Boris^hu/hu^* testes (≥50 tubules scored per mouse; n = 5 biological replicates per genotype). Arrows indicate apoptotic (TUNEL-positive) germ cells. Two-tailed unpaired t-test (p-value < 0.05). (D) Pedigree plots depicting breeding outcomes for *Boris*, *Ctcf*, and combined mutant alleles. Left: previously published crosses demonstrating that *Boris^−/−^* males are subfertile with reduced litter sizes, *Ctcf^+/−^* males are fertile with normal reproductive capacity and that *Ctcf^+/−^* × *Boris^−/−^* double-mutant males are infertile. Upper center and lower right: humanized *Boris^hu/hu^* males show normal fertility, and *Ctcf^+/−^* × *Boris^hu/hu^* double-mutant males also remain fertile. Symbol colors correspond to genotypes listed in the legend. (E) Litter sizes from timed matings. *Boris^hu/hu^* and *Ctcf^+/−^* × *Boris^hu/hu^* crosses produce normal litter sizes, whereas *Ctcf^+/−^* × *Boris^−/−^* crosses are sterile. Two-tailed unpaired t-tests; ns, not significant; **p < 0.001.

Under standard breeding conditions, homozygous humanized males (*Boris^hu/hu^*) displayed normal reproductive parameters at 90 days of age (Fig 3A), the timepoint where *Boris^−/−^* KO mice show clear defects^4,26^. Testis mass, sperm counts, and sperm motility were comparable to wildtype littermates, with no significant differences detected (Fig 3A). Histological analysis revealed normal seminiferous tubule architecture with all stages of germ cell development intact (Fig 3B), contrasting sharply with the testicular atrophy observed in *Boris^−/−^* KO mice at this age^4^. TUNEL staining confirmed no increase in germ cell apoptosis in humanized testes compared to wildtype controls (Fig 3C). Breeding trials showed that *Boris^hu/hu^* males were fully fertile and produced litters of normal size comparable to *Boris^+/+^* controls (Fig 3D-E).

To stringently test functional conservation, we then introduced the humanized *BORIS* allele into the *Ctcf^+/−^* background by double crossing both strains (Fig 3D). We reasoned that the complete infertility of *Ctcf^+/−^ Boris^−/−^* double mutants provides a sensitized genetic background (with a more severe phenotype) to assess whether hBORIS can truly compensate for loss of mBORIS under reduced CTCF dosage (Fig 3D). IHC revealed that hBORIS and CTCF co-localized in nuclei of spermatogonia and primary spermatocytes in humanized testes (Fig 4E), demonstrating proper spatiotemporal expression consistent with published scRNA-seq data from mouse testes^5,36^. This co-localization during the critical window of meiotic initiation supports the capacity for cooperative function between hBORIS and mCTCF. Indeed, *Ctcf^+/−^ Boris^hu/hu^* males were fully fertile and successfully sired multiple litters with typical numbers of live pups (Fig 3E), confirming a successful genetic rescue. This supports that hBORIS not only substitutes for mBORIS function during spermatogenesis but also cooperates effectively with mouse CTCF to support male gametogenesis.

**Figure 4.**
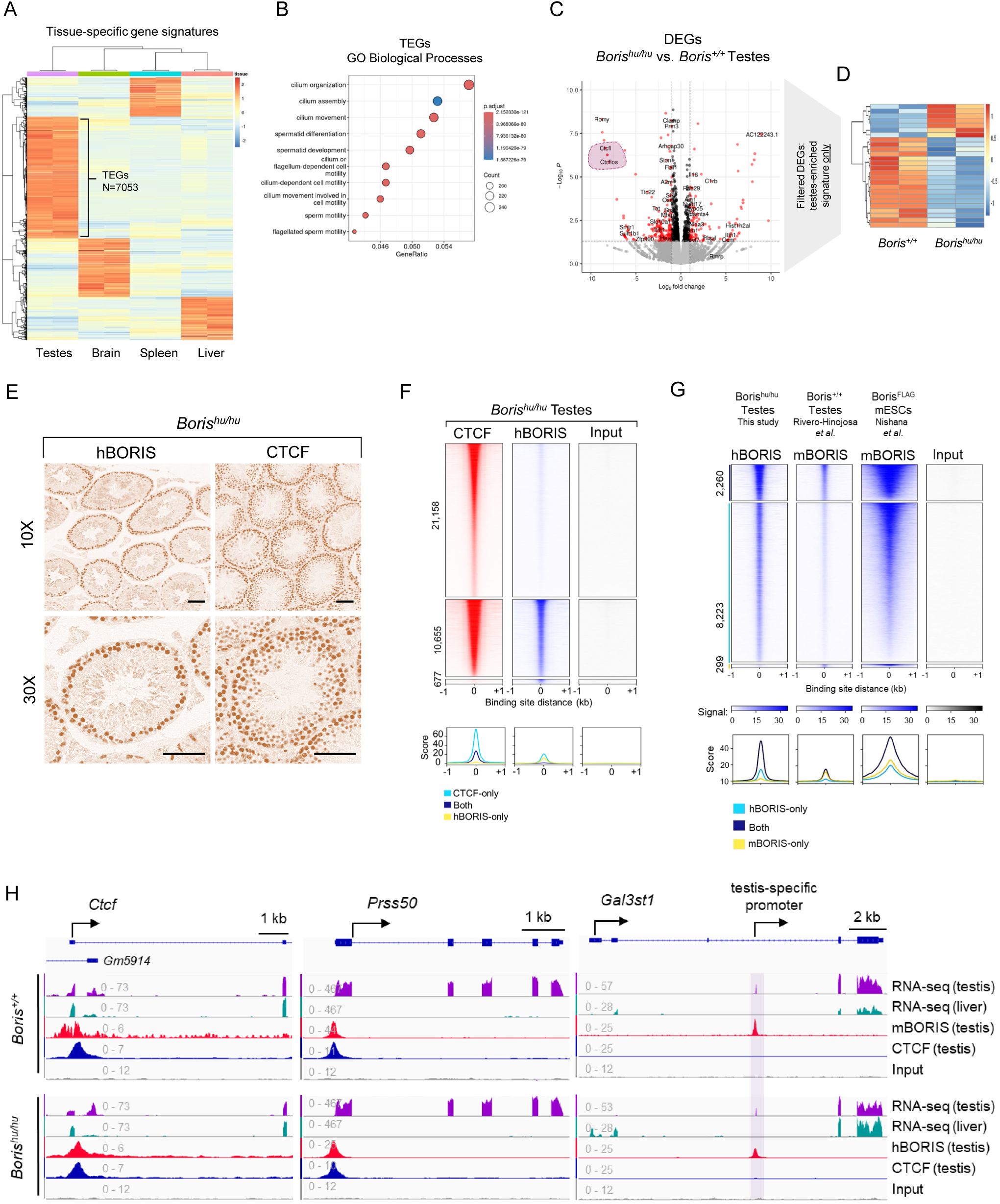
Human BORIS binds conserved germline chromatin regions and maintains core testes transcriptional programs. (A) Heatmap showing tissue-specific RNA-seq expression signatures derived from wild-type testes, brain, liver, and spleen. Differential expression and hierarchical clustering delineate 7,053 testes-enriched genes and 2,215 testes-specific genes (see Fig S4), comprising the germline reference set used in subsequent analyses. (B) GO Biological Process enrichment of the 7,053 testes-enriched genes, highlighting significant enrichment for pathways involved in spermatogenesis, germ cell differentiation, meiotic cell-cycle processes, and cilium/flagellum organization and motility. (C) Differential expression analysis of *Boris^hu/hu^* versus *Boris^+/+^* testes. Volcano plot shows 71 differentially expressed genes (padj < 0.05; |log_2_FC| > 1), highlighting downregulation of the endogenous *Boris/Ctcfl* locus and the antisense transcript *Ctcflos*. (D) Heatmap of testes-enriched differentially expressed genes in *Boris^hu/hu^* testes. Expression patterns show limited perturbation of germline transcriptional programs, with overall preservation of testes-enriched gene signatures. (E) Representative immunohistochemistry for hBORIS and CTCF shows co-localization in early stage germ cells in *Boris^hu/hu^* testes at 10× and 30× magnification. Nuclear DAB signal is observed within seminiferous tubules. Scale bars, 50 μm. (F) Heatmaps of normalized ChIP–seq signal for CTCF and hBORIS in Boris^hu/hu^ testes, centered on factor-specific peak summits (±1 kb) and ranked by intensity. Aggregate profiles (below) show mean coverage across all sites. Peak numbers associated with each cluster are indicated at left. (G) Cross-validation of human BORIS binding sites using previously published mouse BORIS (mBORIS) in *Boris^+/+^* testes ChIP–seq and mBORIS–FLAG ChIP–seq in mESCs. Heatmaps show normalized signal centered on peak summits (±1 kb) for each dataset, confirming that human BORIS in *Boris^hu/hu^* testes occupies orthologous regulatory elements and retains conserved DNA-binding specificity. Aggregate profiles below display mean coverage across the corresponding peak groups. (H) Genome browser snapshots showing representative loci with hBORIS and CTCF ChIP–seq signal in *Boris^hu/hu^* testes. Tracks display normalized ChIP–seq coverage for hBORIS, CTCF, and corresponding RNA-seq profiles from *Boris^+/+^* and *Boris^hu/hu^* testes.

### Human BORIS Shows a High Degree of Functional Conservation in the Murine Germline

Next, we characterized hBORIS function in the humanized mouse germline by defining chromatin occupancy and assessed its impact on transcription. We defined a reference germline transcriptional signature by performing RNA-seq on wildtype testes and compared it to transcriptional signatures from multiple somatic tissues (Fig 4A, Table S2). This approach revealed 7,053 testes-enriched genes (TEGs) (Fig 4B, Table S2), defined as genes showing significantly higher expression in testes than in somatic tissues (Fig S4A). We further defined a subset of 2,215 testes-specific genes (TSGs) with near-exclusive testis expression (Fig S4B, Table S2). Gene ontology analysis of TEGs revealed strong enrichment for germ cell development, spermatid differentiation, meiotic cell cycle, and sperm motility processes (Fig 4B), validating that the TEG signature represented essential spermatogenic transcriptional programs. Functional annotation of somatic transcriptional signatures further validated this tissue-specific approach (Fig S4C, Table S2). The TEG signature described here served as a reference used to define upregulated TEGs (U-TEGs) in somatic contexts later in the study (Fig 5, Table S2).

Previous studies have shown that loss of mBORIS in Boris^−/−^ KO mouse testes results in relatively mild transcriptional defects ^5,26^. To assess the effect of hBORIS expression on the murine spermatogenic program, we compared gene expression in *Boris^hu/hu^* versus *Boris^wt/wt^* testes by RNA-seq (Fig 4C, Table S2). This revealed 71 differentially expressed genes (DEGs; 32 upregulated, 39 downregulated), of which 26 were TEGs and 14 were TSGs (Table S2). As expected, mouse *Boris* (*Ctcfl*) and its antisense lncRNA *Ctcflos* were among the most downregulated genes in *Boris^hu/hu^* testes (Fig 4C, Table S2), validating the replacement of the endogenous locus and confirming that in this context, bulk RNA-seq was suitable to detect biologically relevant changes.

**Figure 5.**
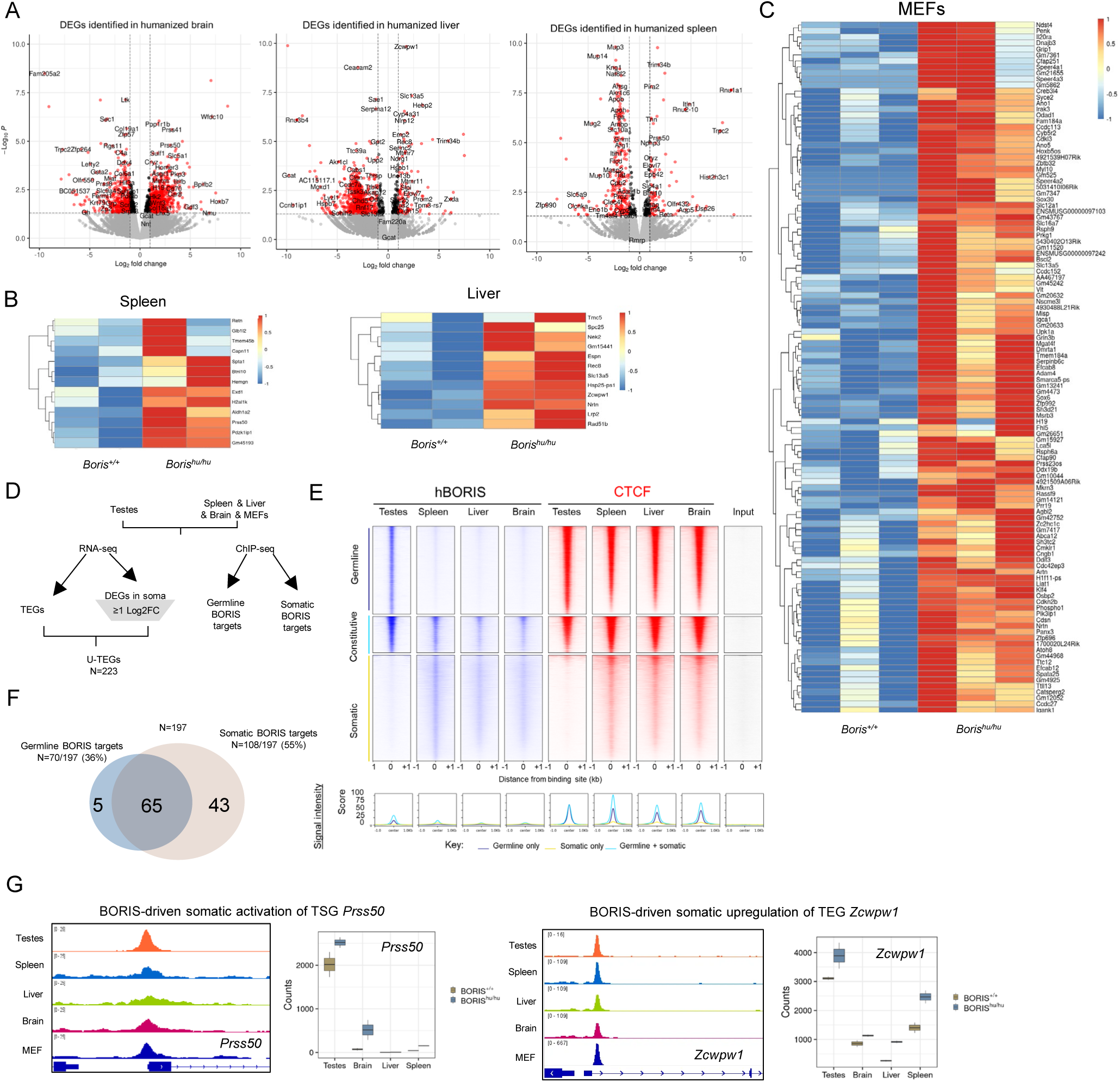
Ectopic activation of germline transcriptional programs and chromatin binding caused by human BORIS expression in somatic tissues. (A) Volcano plots showing differentially expressed genes (DEGs) in *Boris^hu/hu^* brain, liver, and spleen relative to *Boris^+/+^* controls (n = 2 biological replicates). (B) Heatmaps of testes-enriched DEGs upregulated in *Boris^hu/hu^* spleen (n=2), liver (n=2). (C) Heatmaps of testes-enriched DEGs upregulated in *Boris^hu/hu^* MEFs (n=3). See brain in Fig 6C. (D) Schematic illustrating integration of RNA-seq and BORIS ChIP–seq datasets. Upregulated testes-enriched genes in *Boris^hu/hu^* somatic tissues were intersected with germline and somatic BORIS peak sets to define target classes. (E) ChIP–seq heatmaps and aggregate profiles for hBORIS and CTCF across *Boris^hu/hu^* testes, spleen, liver, and brain. Signals are plotted ±1 kb around peak summits for germline-specific, constitutive, and somatic hBORIS-binding classes. Corresponding CTCF ChIP–seq from the same tissues is shown for comparison. Aggregate enrichment plots below display mean signal intensities for germline-only, somatic-only, and shared peak subsets. (F) Venn diagram summarizing the integrated analysis shown in panel D. Of the 223 testes-enriched genes upregulated in *Boris^hu/hu^* somatic tissues, 197 were assessed for BORIS-binding. These comprise germline BORIS targets (70/197; 36%) and somatic BORIS targets (108/197; 55%), with 65 genes shared between categories. (G) Left: Genome browser views of representative testes-enriched genes (e.g., *Prss50, Zcwpw1*) showing ectopic hBORIS binding in *Boris^hu/hu^* somatic tissues. Right: Raw count boxplots for the same loci demonstrating increased expression in *Boris^hu/hu^* brain, liver, and/or spleen. Counts represent two biological replicates per genotype (*Boris^hu/hu^* and *Boris^+/+^*). Additional examples are provided in Fig S8.

In wildtype mice, early-stage germ cells represent the only physiologically normal context where CTCF and mBORIS reported to be co-expressed (Fig 4E), and demonstrate important functional synergy to support spermatogenic transcriptional programming ^5,26^. To assess if this was preserved in the humanized mouse germline, we performed hBORIS and CTCF ChIP-seq in *Boris^hu/hu^* testes (Fig 4F, Fig S5A). hBORIS occupied 10,454 genomic sites (Fig 4F, Fig S5A), binding to 7,164 genes, of which 2,757 were bound directly at the promoter (Fig S5B-C). This represents a four-fold increase in detected peaks compared to previous published mBORIS ChIP-seq data obtained in wildtype mouse testes, when assessed with the same peak calling pipeline^5^ (Fig 4G, Fig S5A). QC comparison between our data and published mBORIS data^37^ strongly suggest the increase in hBORIS peak number is due to better hBORIS-Ab performance and increased sequencing depth (Fig S5D-F). Integration with CTCF ChIP-seq data revealed substantial BORIS-CTCF co-occupancy, with hBORIS detected at 33.5% of all CTCF-bound sites in testes - nearly double the co-occupancy rate previously reported for mBORIS^5^ (17.5%) (Fig 4G). This is consistent with IHC and the functional rescue of *Ctcf^+/−^ Boris^−/−^* infertility (Fig 3) and suggests a greater degree of combined action between hBORIS and CTCF than previously understood. Notably, a subset of hBORIS peaks showed no CTCF co-binding (Fig 4G), suggesting some CTCF-independent functions.

The majority of hBORIS peaks overlapped with previously reported mBORIS peak coordinates^5,37^ (Fig S5A). Furthermore, hBORIS peak summits were strongly enriched for the canonical BORIS DNA-binding motif and showed a high level of sequence conservation across vertebrates (Fig S5G-H). This supports that the additional hBORIS occupancy identified here represents the targeting of orthologous regulatory regions in the mouse genome rather than ectopic binding events. However, a small number of sites bound by mBORIS in previous studies were not detected by hBORIS-specific antibodies, suggesting minor species-specific binding differences (Fig S5A). Notably, these sites exhibited lower motif enrichment and reduced sequence conservation (Fig S5G-H), which could represent mBORIS occupancy at some sites is mediated indirectly through protein-protein interactions – which may not be occupied by hBORIS due to N- and C-terminal sequence divergence.

hBORIS occupancy revealed a broad pattern of genomic distribution, with substantial binding detected at intergenic and intronic regions, as opposed to majority promoter-proximal sites shown by mBORIS (Fig S5B). In addition, 2,757 gene promoters were directly bound by hBORIS (Fig S5C) almost double the number reported for mBORIS^5^ - highlighting both the improved dataset quality and potentially distinct regulatory properties of hBORIS. Gene ontology analysis of hBORIS-bound genes in the humanized mouse germline showed strong enrichment for meiosis, chromosome segregation, DNA recombination, and spermatogenesis pathways (Fig S5C). Integration of ChIP-seq data with TEGs (Fig 4H, Table S2) revealed that 18% of TEGs (1265/7053) and 6% of TSGs (134/2,215) were occupied by hBORIS, supporting that hBORIS specifically regulates core germline transcriptional programs. hBORIS-targeted TEGs serve as molecular markers for ectopic germline programming when examining somatic tissues later in the study.

The modest number of DEGs in *Boris^hu/hu^* testes (Fig 4C, Table S2), combined with normal fertility of *Boris^hu/hu^* mice (Fig 3E), indicates a strong degree of functional conservation between hBORIS and endogenous mBORIS. Only 10 of 71 DEGs identified in *Boris^hu/hu^* testes showed direct hBORIS binding, which may represent subtle species-specific differences in BORIS-mediated gene regulation at a limited subset of genomic loci, rather than widespread functional divergence. Importantly, known mBORIS target genes that are critical for spermatogenesis and fertility—including *Prss50*, *Gal3st1*, and *Ctcf* itself ^4,5,26,27^—showed robust hBORIS occupancy at their promoters with normal expression levels (Fig 4H), demonstrating functional conservation of hBORIS as a transcription factor in the murine germline. Together, these findings establish that hBORIS occupies conserved regulatory elements in the mouse germline, binds extensively at testes-enriched and testes-specific genes through conserved zinc finger-DNA recognition, cooperates with CTCF at chromatin, and maintains expression of core spermatogenesis genes. The high-quality ChIP-seq dataset reveals that functional conservation is mediated by the zinc finger DNA-binding domain, while minor species-specific differences in binding patterns likely reflect N-and C-terminal sequence divergence that affects protein-protein interactions but does not compromise core germline function.

### Ectopic BORIS Expression Drives Germline Gene Activation in Somatic Tissues

Next, we assess whether the ectopic hBORIS expression identified in somatic tissues causes transcriptional deregulation (Fig 2). We performed RNA-seq in somatic tissues from *Boris^hu/hu^* and *Boris^+/+^* mice (Fig 5A), revealing 1,136 DEGs in brain, 839 in liver, and 389 in spleen (Fig 5A, Table S2). The mosaic nature of ectopic hBORIS activation in somatic tissues presented a significant challenge as bulk RNA-seq from heterogeneous tissues captures an average signal across predominantly hBORIS-negative cells, inevitably diluting any resulting transcriptional signature from the hBORIS-expressing minority (Fig S6A). This limitation is compounded by tissue complexity, where diverse cell types each contribute distinct baseline expression patterns that may further obscure the effects of hBORIS.

To address this, we employed two complementary strategies. First, in addition to RNA-seq in somatic tissues, we also performed RNA-seq in clonal hBORIS-positive MEFs (Fig 2F, Fig S6B). Second, we focused our analysis on ectopic activation of testes-enriched genes (TEGs) (Fig 4A, Fig S6B). This approach is biologically justified as TEGs are either silenced or expressed at very low levels in somatic tissues in comparison to testes, making their upregulation in brain, liver, spleen, or MEFs an unambiguous marker of transcriptional deregulation. Unlike housekeeping or ubiquitous genes where expression changes could reflect indirect effects, cell-type composition differences, or technical noise—the ectopic expression of TEGs provided clear evidence of inappropriate gene activation. Applying this stringent filter on DEG lists revealed 93 significantly upregulated testes-enriched genes (U-TEGs) in brain (Fig S6C, Table S2), 12 in liver, 13 in spleen (Fig 5B, Table S2) and 115 in MEFs (Fig 5C, Fig S6E). In total, 223 U-TEGs were identified in the humanized mouse soma (Fig 5D, Table S2).

To determine whether U-TEGs were directly bound by hBORIS, we performed hBORIS ChIP-seq on paired somatic samples used in RNA-seq (Fig 5E, Fig S7F-H). CTCF ChIP-seq was also conducted with peak counts ranging from 55,962 in brain, 57,291 in spleen, 64,244 in liver, and 43,917 in MEFs (Fig 5E, Fig S7A), consistent with CTCF’s ubiquitous role as a chromatin organizer^3^. As expected, the majority of CTCF peaks (40,515) were shared across all tissues (Fig 5E, Fig S7B), with a small fraction showing tissue-specific occupancy. hBORIS binding patterns in somatic tissues predominantly overlapped with CTCF-occupied sites, consistent with co-operative binding observed in the germline (Fig 4F). However, detection sensitivity for hBORIS varied substantially in heterogenous samples with 12,583 peaks identified in spleen, 2,252 in liver, 1,403 in brain; compared to 28,338 in MEFs (Fig 5E, Fig S7B-C). hBORIS peaks in somatic tissues were largely reproducible across different somatic contexts but signal intensities varied greatly in tissues (Fig S7B-C). We anticipate these differences likely reflect the variable proportion of hBORIS-expressing cells in each sample and estimate that the total number of hBORIS binding events identified in somatic tissues are underrepresented due to signal dilution (Fig S7C). hBORIS motif enrichment also varied across somatic tissues suggesting that some ectopic binding events may reflect interactions with other DNA binding proteins (Fig S7D). Peak annotation revealed that hBORIS binding in all somatic tissues was strongly enriched at promoter-proximal regions (±2kb of TSS), contrasting with the broader genomic distribution observed in testes (Fig S7E).

Next, we integrated ChIP-seq and RNA-seq data to assess hBORIS occupancy across U-TEGs identified in the soma of humanized mice (Fig 5D, Table S2). In total, 197/223 U-TEGs had unambiguous genome annotation and sufficient coverage to assess hBORIS occupancy at promoter regions (±5kb of TSS) (Table S2). Systematic scoring of hBORIS ChIP-seq peaks across all five contexts revealed distinct binding patterns (Fig 5F). Among the 197 genes analyzed, 70 genes (36%) showed detectable hBORIS binding in testes (Table S2), identifying these as established hBORIS regulatory targets in their native context (Fig 5F). Of these germline hBORIS targets, 65 (93%) also showed detectable BORIS occupancy in at least one somatic context, while 5 showed germline binding only (Fig 5F, Table S2)—likely reflecting somatic binding below detection threshold in heterogeneous tissues. An additional 43 genes showed detectable somatic hBORIS occupancy without corresponding germline peaks, highlighting examples of ectopic hBORIS binding and potentially reflecting context-specific chromatin accessibility between cell types. In total, 108 upregulated genes (55%) showed detectable hBORIS occupancy in at least one somatic context (Fig 5F, Table S2), while 84 genes (43%) showed no detectable hBORIS binding in any context examined (Table S2), suggesting these represent indirect targets, enhancer-regulated genes or genes activated through cooperative effects with other hBORIS-induced transcription factors. This strong concordance demonstrates that ectopic BORIS retains its native DNA-binding specificity and reactivates established germline regulatory programs when expressed in somatic cells.

Examples of ectopic hBORIS-driven causing upregulation of TEGs are shown in Fig 5G. *Prss50,* crucial for spermatogenesis, showed complete reactivation from a silenced state (Fig 5G). *Prss50* showed robust hBORIS occupancy at its promoter across multiple somatic contexts and showed exclusive activation in brain and spleen of *Boris^hu/hu^* mice (Fig 5G). In addition, *Zcwpw1*, a histone reader involved in meiotic recombination^38^, also showed robust hBORIS binding at its promoter all somatic contexts with coordinated upregulation in all hBORIS-expressing tissues examined (Fig 5G). Additional TEGs showing hBORIS binding and ectopic activation included *Exd1* (exonuclease involved in DNA repair), *Steap1* (metalloreductase), and *Xrcc2* (DNA recombination factor) (Fig S8), reinforcing that germline developmental genes are reactivated by ectopic hBORIS expression.

Beyond the targeted analysis of TEGs, we also observed several upregulated cancer-associated genes in humanized somatic tissues (Fig S8). In liver, *Rec8* (subunit of meiotic cohesin), *Spon2* (extracellular matrix protein), and *Ndrg1* (metastasis suppressor) all showed activation. In brain, *Wfdc10* (WAP domain protein), *Epb41l1* (cytoskeletal regulator), and *Prss41* (serine protease) were significantly upregulated. In spleen, *Rnu1a1* (small nuclear RNA component) and *Serpinc1* (serine protease inhibitor) also showed elevated expression. *Hebp2* (heme-binding protein implicated in microtubule dynamics and cancer pathways^39^ was activated in spleen with corresponding hBORIS occupancy at its promoter. While these cancer-associated genes were not uniformly testes-enriched, their ectopic activation in hBORIS-expressing tissues demonstrates that hBORIS-driven transcriptional disruption does extend beyond germline-specific genes to include broader transcriptional destabilization that includes oncogenic targets.

### Ectopic hBORIS Expression Increases Cancer Susceptibility

Having established that hBORIS drives inappropriate transcriptional reprogramming in somatic tissues (Fig 5), we investigated whether this translated to pathological consequences *in vivo*. Given that hBORIS is a designated human CTA, we hypothesized that somatic hBORIS expression might increase cancer susceptibility in *Boris^hu/hu^* mice. To assess long-term health consequences, we monitored cohorts of *Boris^hu/hu^* (n=88) and *Boris^wt/wt^* (n=56) littermates throughout their natural lifespan under identical specific pathogen-free housing conditions with standardized diet and veterinary care. Animals underwent routine health evaluations with endpoint euthanasia upon development of clinical signs of morbidity, including hunched posture, labored breathing, significant weight loss or palpable masses (Fig 6A).

**Figure 6.**
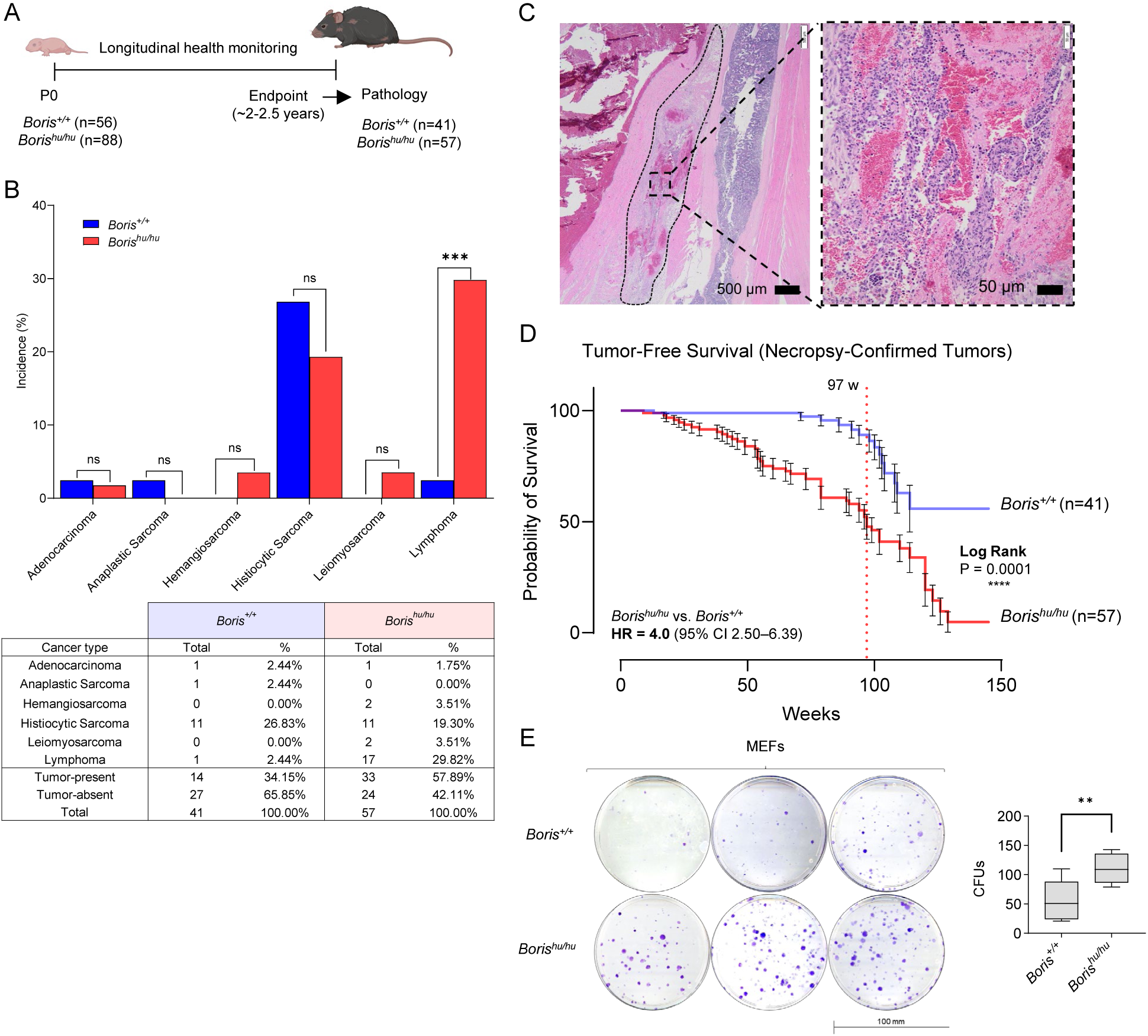
Somatic hBORIS expression reduces survival and increases cancer susceptibility. (A) Study design illustrating longitudinal monitoring of *Boris^+/+^* (n = 56) and *Boris^hu/hu^* (n = 88) mice from P0 through natural endpoint (~2–2.5 yr), followed by pathological assessment of surviving cohorts (*Boris^+/+^*, n = 41; *Boris^hu/hu^*, n = 57). (B) Tumor incidence in *Boris^+/+^* and *Boris^hu/hu^* (n = 57) mice at endpoint. Bar plots show the proportion of animals developing each tumor type within each genotype cohort, with corresponding counts summarized in the table below. No significant differences were observed for adenocarcinoma, anaplastic sarcoma, hemangiosarcoma, histiocytic sarcoma, or leiomyosarcoma (Fisher’s exact test, ns). Lymphoma incidence was significantly elevated in *Boris^hu/hu^* mice compared with *Boris^+/+^* (***P < 0.001, Fisher’s exact test). (C) Representative image of rare hemangiosarcoma in *Boris^hu/hu^* mice showing atypical endothelial cells lining blood-filled spaces. Scale bars, 50 μm. (D) Tumor-free survival in *Boris^+/+^* (n = 41) and *Boris^hu/hu^* (n = 57) mice based on necropsy-confirmed tumor diagnoses. Kaplan–Meier curves show a significantly earlier onset of tumor development in *Boris^hu/hu^* animals compared with *Boris^+/+^*, with a hazard ratio of 4.0 (95% CI, 2.50–6.39). The dotted line marks median tumor-free survival (97 weeks). Survival distributions were compared by log-rank test (P = 0.0001). (E) Clonogenic assay showing enhanced colony-forming capacity in hBORIS-positive MEFs (see Fig 2). Above: Representative crystal violet-stained plates from *Boris^+/+^* and *Boris^hu/hu^* MEFs (3 independent clones plus 3 technical replicates per genotype). Below: Quantification of colony number. Data shown as mean ± SEM; **p<0.01, unpaired t-test.

Kaplan-Meier analysis of all-cause mortality across the complete cohort revealed significantly reduced overall survival in *Boris^hu/hu^* mice compared to *Boris^+/+^* controls (Fig S9A). Median overall survival was 90 weeks in *Boris^hu/hu^* mice versus 102 weeks in *Boris^+/+^* controls (log-rank test: χ^2^=7.33, df=1, p=0.007, hazard ratio =1.58, 95% confidence interval: 1.13–2.20). Comprehensive necropsy with systematic multi-organ examination and histopathological analysis was performed on all animals euthanized for clinical deterioration with adequate tissue preservation. Any euthanized animals found in a state of tissue autolysis upon examination were excluded from the study. Board-certified veterinary pathologists rendered all diagnoses, identifying an elevated incidence of cancer in 33/57 (57.89%) *Boris^hu/hu^* mice, compared to only 14/41 (34.15%) *Boris^+/+^* littermates, representing a 1.6-fold increase (χ^2^=4.12, df=1, p=0.042) in humanized mice (Fig 6B). All neoplastic and non-neoplastic pathological findings and statistical analyses are provided in Table S3.

Hematologic malignancies (histiocytic sarcoma and lymphoma) predominated in both genotypes, affecting 12/41 (29%) wildtype and 28/53 (49%) humanized mice (Fig 6B). Histiocytic sarcoma, occurred in 11/41 mice (26%), and similarly, 11/57 (19%) humanized mice—consistent with previously published incidence rates in the C57BL/6 strain ^40^ (Fig 6B, Fig S9B-D). While rates of histiocytic sarcoma did not differ between genotypes (Fig 6B), humanized mice exhibited a profound shift toward lymphoid malignancies, with lymphomas in 17/53 animals (30%) compared to only 1/41 (2.4%) in wildtype controls (Fisher’s exact test, p=0.0004) (Fig 6B, Fig S9B-D). The lymphomas arising in 12/17 (70%) of *Boris^hu/hu^* mice were high-grade and multifocal (two or more anatomical sites) - largely involving spleen, lymph nodes, liver, thymus, and bone marrow (Fig 6B, Fig S9E-G, Table S3). Beyond hematological malignancies, some *Boris^hu/hu^* mice developed additional malignant tumors including leiomyosarcoma (n=2) and highly aggressive hemangiosarcoma (n=2), (Fig 6C). There tumors were not observed in *Boris^+/+^* controls. While case numbers limited statistical analysis, the exclusive emergence of these rare and aggressive tumors suggests ectopic hBORIS expression confers broad multi-lineage cancer susceptibility beyond hematopoietic cells.

Among mice with diagnosed cancer, median tumor-free survival in *Boris^+/+^* mice was not reached as >50% of mice remained tumor free, whereas median-tumor free survival in *Boris^hu/hu^* mice was 97 weeks (log-rank test: χ^2^=25.80, df=1, p<0.0001, HR=4.00, 95% CI: 2.50–6.39) (Fig 6D). The Gehan-Breslow-Wilcoxon test, specifically used to test for differences in survival distributions (with more weight given to early events), showed highly significant divergence (χ^2^=28.44, p<0.0001), demonstrating that tumor-bearing humanized mice reach clinical endpoints much earlier in life than tumor-bearing wildtype mice (Table S3). Critically, the hazard ratio for confirmed tumor presence (HR=4.00) substantially exceeded that for all-cause mortality (HR=1.58), demonstrating that malignancy—not generalized systemic pathology—drives excess mortality in humanized mice (Fig 6D). Strikingly, no wildtype mice succumbed to tumors before 60 weeks of age, whereas 18/57 (31%) of humanized mice presented with morbid malignancy before this timepoint (Fisher’s exact test, p=0.001). This early-onset phenotype indicates that hBORIS expression fundamentally alters cancer biology, accelerating tumorigenesis rather than merely increasing susceptibility to age-associated malignancy.

This cancer susceptibility phenotype reflects incomplete developmental silencing of the humanized *BORIS* allele, resulting in persistent hBORIS expression and inappropriate activation of germline transcriptional programs. To determine whether this directly confers transformation potential at the cellular level, we assessed proliferative capacity in MEFs. Clonogenic assays demonstrated that hBORIS-positive MEFs exhibited significantly enhanced colony formation capacity versus wildtype MEFs without BORIS expression (Fig 2F, Fig 6E). This increased clonogenic potential – a hallmark of transformation - provides direct evidence that hBORIS expression enhances proliferative capacity and resistance to contact inhibition, establishing a mechanistic link between ectopic hBORIS-driven germline gene activation and organismal cancer susceptibility.

Together, these findings demonstrate that ectopic hBORIS expression—arising from incomplete developmental silencing of the humanized *BORIS* allele—increases early-onset and overall cancer incidence, shifts hematologic malignancy spectrum toward aggressive lymphomas, and expands solid tumor diversity. These results establish a direct link between regulatory divergence at the *BORIS* locus (Fig 2), inappropriate hBORIS-driven germline gene activation in somatic tissues (Fig 5), and cancer predisposition (Fig 6) - validating the cancer-testis antigen model of hBORIS function in tumorigenesis.

## Discussion

To define the functions of human BORIS *in vivo*, we generated a genetically humanized BORIS mouse model which retained human specific cis-regulatory elements, designed to support endogenous regulatory control of the human locus. Following humanization of the *Boris* locus, human *BORIS* was expressed from the canonical promoter, in the appropriate spatio-temporal pattern in the germline^4,5^, rendering our strategy to include upstream and downstream regulatory elements successful.

Mouse and human spermatogenesis differ substantially in timing, gene expression programs, and regulatory requirements^41^, therefore we first investigated whether hBORIS could functionally substitute for mBORIS in the mouse germline, despite its sequence divergence. Integrated ChIP-seq and RNA-seq analyses in testes revealed that the central zinc finger domain is sufficiently conserved to mediate hBORIS DNA binding at conserved mBORIS targets genomic sites and support transcription of key mBORIS-dependent fertility genes—including *Prss50*, *Gal3st1*, and *Ctcf*^4,5,26,27^. Furthermore, our ChIP-seq data revealed thousands more hBORIS occupied sites in the germline than previously profiled by mBORIS ChIP-seq, with extensive CTCF co-occupancy, suggesting a broader combined action of CTCF and BORIS in spermatogenic programming than previously understood^5^. Beyond known mBORIS targets, the transcription of most testes-specific genes in the humanized germline were minimally affected. Despite substantial sequence divergence in the N- and C-terminal domains^1^ our data supports that hBORIS retains a high degree of functional conservation with mBORIS – although species-specific functions are yet to be elucidated^25,28^. The substitution of mBORIS with hBORIS fully rescued both the previously reported baseline subfertility of *Boris* knockout mice and the complete sterility of *Ctcf^+/−^ Boris^−/−^* double mutants^4,26,42^. These data effectively demonstrate how hBORIS can integrate into the murine germline regulatory network and support transcriptional regulation essential for spermatogenesis, and underscores a selective pressure to maintain core BORIS function across species.

During routine characterization of the humanized mouse model, we detected mosaic hBORIS expression, restricted to a limited number of cells in somatic tissues, presenting both challenges and opportunities for exploring its role outside the germline, in line with its classification as a cancer-testis antigen^1,16,43^. Previous reports described that constitutive *Boris* expression during embryonic development results in severe vascular defects and neonatal lethality^44^. Not only did we detect hBORIS expression in adult tissues, but hBORIS was also robustly expressed in mESCs and MEFs, suggesting it may be expressed throughout embryonic and developmental stages. In contrast to previous reports, we observed normal viability and no developmental abnormalities (Fig 3, Table S3), highlighting that *BORIS* can be tolerated in subsets of cells during development without causing developmental lethality. Extensive molecular profiling of humanized mice transcriptome, across several somatic contexts, revealed >200 ectopically upregulated testes-enriched genes (U-TEGs), including meiotic regulators (*Mei4*)^45^, germline-specific genes (*Prss50, Zwcwp1*)^27,46^, developmental genes (Aldh1a2, Rec8, Sox6, Wnt3/6) and other cancer-testis antigens (Prss41, Syce2, Sycp1)^47–49^. The detection of a robust germline gene activation signature in the soma, despite sample heterogeneity (hBORIS protein detected in relatively few somatic cells) indicates strong transcriptional effects within hBORIS-positive cells. We confirmed hBORIS occupancy at the majority of U-TEGs in somatic tissues and verified that these were also direct hBORIS targets in the germline, demonstrating that hBORIS reactivates its native germline regulatory program when aberrantly expressed. Several U-TEGs genes showed no detectable hBORIS occupancy in promoter-proximal regions (±5kb). Our analysis focused on promoter-proximal regions, potentially missing distal enhancer binding events. BORIS and CTCF have both overlapping and distinct regulatory functions^50^, with BORIS capable of functioning as a transcription factor at promoters while also potentially disrupting CTCF-mediated enhancer-promoter loops, which are essential for normal cellular function^7,51^. This finding places hBORIS ectopic activity within the context of germline program reactivation in cancer. Increasing numbers of studies document germline-specific transcriptional as a phenomenon linked to cellular dedifferentiation and increased plasticity associated with cancer^52–55^. For example, Mei4 activation induces genome instability through inappropriate DNA double-strand break formation and recombination in non-meiotic contexts^45^. Germline developmental programs also confer proliferative capacity and dedifferentiation phenotypes that support tumor initiation and progression. As shown in our data, hBORIS can induce activation of additional CTAs (*Prss41, Syce2, Sycp1*), which may trigger a cascade of genome instability and destabilized cell identity, as many CTAs themselves encode transcriptional regulators or chromatin modifiers^56^.

In this model, ectopic hBORIS expression significantly increased cancer incidence and reduced overall survival, with a substantial fraction of humanized mice developing morbid malignancies early in life. This early tumor-associated mortality indicates that hBORIS accelerates tumorigenesis rather than simply increasing susceptibility to age-associated cancers, supporting a direct role for aberrant germline program activation in shaping somatic cancer risk^52–55^. In addition to increased cancer burden, hBORIS expression was associated with a pronounced shift in tumor spectrum. While histiocytic sarcoma—typical of the C57BL/6 background—occurred at comparable frequencies across genotypes, humanized mice exhibited a striking enrichment for aggressive lymphomas. This shift suggests cell-type–specific oncogenic consequences of ectopic germline programming, potentially reflecting heightened vulnerability of lymphoid compartments. Although the roles of BORIS and CTCF in immune cell regulation remain incompletely defined^57^, the preferential emergence of lymphoid malignancies suggests that lymphoid compartments may be particularly sensitive to disruption of CTCF-associated chromatin regulation by ectopic hBORIS expression. Consistent with this, aberrant BORIS expression has been linked to human lymphoid cancers^7^. Collectively, these data demonstrate that ectopic hBORIS expression actively promotes tumorigenesis in vivo, providing direct evidence that BORIS activation can function as an early oncogenic event rather than a passive marker of dedifferentiation. Accordingly, hBORIS emerges as an oncogenic driver that increases cancer susceptibility, although whether full malignant transformation requires cooperating mutations remains unresolved. Increased clonogenic capacity of hBORIS-positive MEFs supports a cell-autonomous contribution to transformation, although non–cell-autonomous effects cannot be excluded.

The mosaic hBORIS expression pattern observed in this model is consistent with a mosaic epimutation^58^, and closely resembles heterogeneous CTA expression in human tumors^59^, thus providing a physiologically relevant platform for further understanding CTA activation and evaluating CTA-targeted therapies^33^. We propose that somatic hBORIS expression arises from species-specific regulatory incompatibility between the human locus and the mouse epigenetic environment, consistent with the greater regulatory complexity of the human BORIS locus^29^. Although the human BORIS promoter is normally methylated and silenced in somatic tissues^2^, CpG-rich DNA intrinsically resists DNA methylation and requires sustained repression^60,61^, which may be incompletely maintained in a mouse epigenetic environment where species-specific silencing factors are absent or less efficient^62^. The model also enables investigation of CTA function in both the germline, where hBORIS is essential, and in cancer, where it drives transformation, allowing assessment of therapeutic window—an important consideration for CTA-targeted therapies, particularly in cancers affecting individuals of reproductive age, such as ovarian cancer where BORIS is frequently activated^16^. More broadly, our findings not only highlight challenges in modeling genes with rapidly evolving regulatory elements across species^63^, but illustrate how regulatory divergence between species can also inadvertently lead to pathological gene activation that mimics clinically relevant epimutations, that would otherwise be difficult to generate through traditional overexpression approaches.

Here, we established a genetically humanized BORIS mouse model and show that, while hBORIS retains core germline transcription factor function sufficient to support normal spermatogenesis, incomplete developmental silencing of the humanized locus results in persistent mosaic somatic expression that drives ectopic germline gene activation and increased cancer susceptibility in vivo. By preserving endogenous regulatory control in the germline and stochastically escaping silencing in the soma, this model provides a physiologically relevant system with dual utility to study human cancer-testis antigen biology across fertility and cancer contexts.

## Methods

### Generation of Humanized BORIS Mouse Model

The humanized BORIS/BORIS mouse model was generated by Leveragen, Inc. (Boston, USA) using a recombinase-mediated genomic replacement (RMGR) strategy. This method involved the targeted replacement of the endogenous mouse BORIS locus with the human BORIS gene. The process included the use of CRISPR/Cas9 technology to introduce specific docking sites and the subsequent insertion of the human transgene. The human BORIS (hBORIS) sequence was obtained from the a BAC clone. This BAC clone includes a large genomic fragment encompassing the hBORIS gene and its regulatory regions. The hBORIS sequence was confirmed and annotated according to the NCBI Reference Sequence database (RefSeq: NM_080618). All procedures, including vector construction and genetic modifications were carried out under proprietary and confidential protocols developed by Leveragen, Inc.

### Long-Read Sequencing

Long-read sequencing was performed by Leveragen using PacBio SMRT technology to verify the structural integrity of the human *BORIS* insertion. Reads were aligned to the reference genome using pbmm2, and variants were called with GATK HaplotypeCaller to identify structural alterations at the insertion site. Variant effects were annotated using Ensembl Variant Effect Predictor (VEP), and alignments were visually inspected in Integrative Genomics Viewer (IGV) to confirm insertion fidelity and exclude unintended genomic modifications.

### Genotyping PCR and Sequencing

Integration of the humanized BORIS allele was confirmed by PCR followed by sequencing. Genomic DNA was extracted from ear or tail biopsies using a standard lysis buffer protocol. PCR was performed in 25 µL reactions containing 10–50 ng genomic DNA, 0.2 µM of each primer, 200 µM dNTPs, 1× PCR buffer, 1.5 mM MgCl_2_, and 0.5 U Taq polymerase (Invitrogen Platinum Taq). Thermal cycling conditions consisted of an initial denaturation at 95 °C for 5 min; 35 cycles of 95 °C for 30 s, 60 °C for 30 s, and 72 °C for 1 min; followed by a final extension at 72 °C for 7 min. Allele-specific primers were used to selectively amplify the human BORIS insertion (forward: 5′-ACGTACTGAGGAGGCTTCCA-3′; reverse: 5′-GTGGGATCCTAGTGGCACTA-3′), with additional primers used to detect the endogenous mouse Boris allele. PCR products were resolved on 1.5% agarose gels stained with ethidium bromide or SYBR Safe DNA Gel Stain (Thermo Fisher Scientific) and visualized under UV illumination. Expected amplicon sizes were ~500 bp for the humanized BORIS allele and ~350 bp for the wild-type allele.

### Ethical Compliance, Animal Housing and Husbandry

All animal procedures were approved by the NIH and conducted in accordance with institutional guidelines (protocol LIG-15). C57BL/6J mice were used as the background strain for all genetic modifications and were housed under specific pathogen-free conditions with a 12-h light/dark cycle at 22 ± 2 °C, with ad libitum access to standard chow and water. Animals were monitored daily for health and welfare, and all efforts were made to minimize suffering.

### Breeding and Maintenance of Humanized BORIS Mouse Line

To establish a stable homozygous humanized *BORIS* line, heterozygous founder mice carrying one human *BORIS* allele and one endogenous mouse *Boris* allele were generated by targeted CRISPR-mediated insertion. Heterozygous founders were bred to produce F1 offspring, which were subsequently intercrossed to generate F2 progeny containing homozygous humanized, heterozygous, and wild-type genotypes in Mendelian ratios. Genotyping was performed at each generation using PCR on genomic DNA isolated from ear or tail biopsies. Allele-specific primers were used to distinguish the humanized *BORIS* allele (971 bp) from the endogenous mouse *Boris* allele (875 bp). Homozygous humanized mice were selected for experimental studies, while heterozygous mice were maintained for colony propagation.

### Euthanasia and Tissue Collection

Mice were euthanized in accordance with IACUC-approved institutional guidelines using CO_2_ inhalation followed by cervical dislocation. Tissues were collected immediately to preserve RNA and protein integrity. Testis, spleen, brain, and liver were rapidly dissected, rinsed in cold phosphate-buffered saline (PBS), and processed according to downstream applications. For molecular analyses, tissues were either flash-frozen in liquid nitrogen and stored at −80 °C or preserved in RNAlater (Thermo Fisher Scientific). For histological analyses, tissues were fixed in 10% neutral-buffered formalin for 24 h and transferred to 70% ethanol prior to paraffin embedding.

### Tissue Processing

Following euthanasia, testis, spleen, brain, and liver were rapidly dissected, rinsed in cold phosphate-buffered saline (PBS), and fixed in 10% neutral-buffered formalin for 24 h at room temperature. Fixed tissues were submitted to HistoServ, Inc. (Germantown, MD) for paraffin embedding, sectioning, and downstream histological and immunohistochemical analyses.

### Hematoxylin and Eosin Staining

Paraffin-embedded testicular sections (5 µm) were prepared and stained with hematoxylin and eosin by HistoServ, Inc. using standard histological procedures. Stained sections were examined by light microscopy to assess testicular morphology, including seminiferous tubule architecture, germ cell organization, and interstitial structure.

### Immunohistochemistry

Immunohistochemistry was performed by HistoServ, Inc. on paraffin-embedded tissue sections using standard protocols. Sections were stained with an antibody against human BORIS (Abcam; EP12204) to assess expression and localization of the humanized BORIS transgene in testicular and somatic tissues. Brightfield images were acquired by the service provider and analyzed using QuPath software to evaluate staining intensity and cellular localization.

### TUNEL Assay for Apoptosis Detection

Apoptotic cells in testicular tissue sections were detected by terminal deoxynucleotidyl transferase dUTP nick end labeling (TUNEL) staining performed by HistoServ, Inc. using established protocols. TUNEL-positive cells were quantified relative to total nuclei, and the apoptotic index was calculated as the percentage of TUNEL-positive cells per total cells counted.

### Computer-Assisted Sperm Analysis (CASA)

Sperm analysis was performed by the NIH Mouse Genomics Unit using computer-assisted sperm analysis (CASA). Male mice (90 days old) were euthanized, and cauda epididymides were dissected and incubated in pre-warmed culture medium (37 °C) to allow sperm to swim out. Sperm concentration and motility parameters were quantified using an automated CASA system according to the facility’s standard operating procedures. Measured parameters included sperm concentration (million sperm/mL), total motility, and progressive motility. All analyses were conducted at 37 °C.

### Long-Term Survival and Pathology Analysis

For long-term survival studies, mice were aged under specific pathogen-free (SPF) conditions with ad libitum access to standard chow and water. Animals were monitored daily for clinical signs of morbidity, including hunched posture, ruffled fur, labored breathing, reduced activity, significant weight loss, palpable masses, or other signs of distress. Animals meeting IACUC-approved humane endpoint criteria were evaluated by veterinary staff and euthanized accordingly.

At euthanasia, date and clinical presentation were recorded, followed by immediate comprehensive necropsy. Major organs, including lymph nodes, spleen, liver, lungs, kidneys, heart, brain, thymus, bone marrow, gastrointestinal tract, and reproductive organs, were systematically examined. Tissues were fixed in 10% neutral-buffered formalin, paraffin-embedded, sectioned, and stained with hematoxylin and eosin (H&E). All histological sections were evaluated by board-certified veterinary pathologists, and diagnoses were rendered using standardized nomenclature. Tumors were classified by tissue of origin, histological type, and grade where applicable.

### Survival and Statistical Analysis

Overall survival and tumor-free survival were analyzed using Kaplan–Meier curves, with statistical significance assessed by log-rank (Mantel–Cox) tests. Tumor-free survival analyses censored animals euthanized for non-tumor-related causes. The Gehan–Breslow–Wilcoxon test was used to assess differences in survival distributions with emphasis on early events. Hazard ratios (HR) and 95% confidence intervals (CI) were calculated using Cox proportional hazards regression. Tumor incidence between genotypes was compared using Fisher’s exact test or chi-square tests, as appropriate. All statistical analyses were performed using GraphPad Prism v9.0, with p < 0.05 considered statistically significant.

### qPCR

Total RNA was extracted from mouse tissues, such as testis and spleen, using the Direct-Zol RNA Miniprep Kit (Zymo) according to the manufacturer’s protocol. RNA concentration and purity were assessed with a NanoDrop spectrophotometer, ensuring an A260/A280 ratio between 1.8 and 2.1. For cDNA synthesis, 1 µg of total RNA was reverse transcribed using the SuperScript VILO cDNA Synthesis Kit (Thermo Fisher Scientific) in a reaction volume of 20 µL, following the manufacturer’s instructions. cDNA was diluted at least 1:50. Quantitative PCR was carried out on a Applied Biosystems™ QuantStudio™ 5 (Applied Biosystems) thermal cycler. Each qPCR reaction included 1 µL of cDNA, 0.2 µM of each primer, and 1X Power SYBR™ Green PCR Master Mix (Applied Biosystems) in a total volume of 10 µL. Thermal cycling conditions were set to an initial denaturation at 95°C for 10 minutes, followed by 40 cycles of 95°C for 15 seconds and 60°C for 1 minute. Relative expression levels were calculated using the 2^-ΔΔCt method.

### Western Blot

For Western blot analysis, protein was extracted from tissues by homogenization in RIPA buffer (Thermo Fisher Scientific) containing protease inhibitors (Sigma-Aldrich). 20 µL of total protein per sample was loaded onto a 10% SDS-PAGE gel. Proteins were separated by electrophoresis and then transferred onto a PVDF membrane (Millipore) at 100V for 1 hour. The membrane was blocked with 5% non-fat milk in PBS-T (0.1% Tween-20) for 1 hour at room temperature and incubated overnight at 4°C with a primary antibody against human BORIS (1:1000 dilution; Abcam; EP12204). Following primary antibody incubation, the membrane was washed with PBS-T and incubated with HRP-conjugated secondary antibodies (1:5000 dilution) for 1 hour at room temperature. Protein bands were visualized using SuperSignal™ West Pico PLUS Chemiluminescent Substrate (Thermo Scientific) and detected using a ChemiDoc Imaging System (Bio-Rad).

### Chromatin Immunoprecipitation

Tissue samples were crosslinked by adding formaldehyde to a final concentration of 1% and incubating at room temperature for 10 minutes. Crosslinking was quenched by adding glycine to a final concentration of 125 mM and incubating for an additional 5 minutes. Tissues were then washed with cold phosphate-buffered saline (PBS) and subsequently homogenized. Following crosslinking, chromatin was isolated and subjected to shearing to produce DNA fragments averaging 200–500 bp in length. Sonication was performed using a Bioruptor (Diagenode) with settings of 30 seconds on, 30 seconds off, at high power, for 10–15 cycles, with samples kept on ice to prevent overheating. For immunoprecipitation, sheared chromatin was incubated overnight at 4°C with 5 µg of antibodies specific to human BORIS (lab derived monoclonal^13^) or CTCF (Santa Cruz; sc-271514) to capture the target protein-DNA complexes. Antibody-chromatin complexes were then bound to DiaMag protein A-coated magnetic beads (Diagenode) with gentle rotation for 2 hours at 4°C. After binding, beads were washed with increasing stringency using low-salt, high-salt, and lithium chloride buffers, followed by a final wash with TE buffer to remove any non-specific interactions. Following the immunoprecipitation and wash steps, protein-DNA crosslinks were reversed by incubating the samples at 65°C for 4 hours in the presence of proteinase K. DNA was purified using ChIP DNA Clean & Concentrator Kit (Zymo), following the manufacturer’s instructions.

### ChIP-seq

ChIP DNA libraries were prepared using the NEBNext Ultra II DNA Library Prep Kit (New England BioLabs) according to the manufacturer’s instructions. Briefly, 10–20 ng of ChIP DNA was end-repaired, A-tailed, adapter-ligated, and PCR-amplified using Illumina-compatible adapters. Library quality and size distribution were assessed using an Agilent 2100 Bioanalyzer (High Sensitivity DNA Kit), and concentrations were quantified using a Qubit Fluorometer (Thermo Fisher Scientific). Libraries with mean fragment sizes of 200–500 bp were selected for sequencing. Sequencing was performed on an Illumina NovaSeq platform with paired-end reads. Reads were aligned to the mouse reference genome (mm10) using Bowtie2 (v2.4.2)^64^. Peak calling was performed with MACS2^65^ (v2.2.7) using default parameters. Peak annotation was carried out using ChIPseeker^66^ (v1.24.0) to classify binding sites by genomic context.

### RNA-seq

Total RNA was extracted from mouse tissues, including testis and somatic organs, using TRIzol reagent (Thermo Fisher Scientific) followed by purification with the Direct-zol RNA MiniPrep Kit (Zymo Research), according to the manufacturers’ instructions. RNA quality and integrity were assessed using an Agilent Bioanalyzer, and only samples with high RNA integrity numbers (RIN) and A260/A280 ratios between 1.8–2.1 were used for library preparation. RNA was stored at −80 °C until processing. For library preparation, polyadenylated mRNA was isolated from 1 µg of total RNA using the NEBNext Poly(A) mRNA Magnetic Isolation Module (New England BioLabs) and converted into sequencing libraries using the NEBNext Ultra II RNA Library Prep Kit for Illumina. Libraries were fragmented, reverse transcribed, end-repaired, adapter-ligated, and amplified by limited-cycle PCR. Library size distribution (200–400 bp) was confirmed using an Agilent Bioanalyzer. Sequencing was performed on an Illumina NovaSeq platform with paired-end reads. RNA-seq reads were quality-checked using FastQC (v0.11.9) and trimmed using Trimmomatic^67,68^ (v0.39). Reads were aligned to the mouse reference genome (mm10) using HISAT2^69^ (v2.2.1). Gene-level counts were generated using featureCounts (v2.0.1), and differential gene expression analysis was performed using DESeq2^70^ (v1.30.1) in R. Genes with an adjusted p-value < 0.05 (Benjamini–Hochberg correction) and an absolute log_2_ fold change >1 were considered significantly differentially expressed.

### Differentiation of Mouse Embryonic Stem Cells

Mouse embryonic stem cells (mESCs) were cultured using the ESGRO-2i medium system for feeder-free, serum-free culture of mouse ES cells (Millipore, SF001-500) according to the manufacturer’s instructions. Cells were maintained on gelatin-coated tissue culture plates at 37°C in a humidified atmosphere containing 5% CO_2_ and passaged every 2-3 days using 0.05% trypsin-EDTA (Gibco). For retinoic acid (RA)-induced differentiation experiments, mESCs were plated at a density of 2 × 10^4^ cells/cm^2^ on gelatin-coated plates. Retinoic acid (Sigma-Aldrich) was added to the culture medium at a final concentration of 1 μM. Medium was replaced daily with fresh RA-containing medium. Cells were harvested at 0, 24, 48, 72, and 96 hours post-RA treatment for RNA extraction and gene expression analysis. Differentiation was monitored by assessing the downregulation of pluripotency markers (*Oct4*, *Nanog*) and the expression of differentiation markers by qRT-PCR.

### Generation of Mouse Embryonic Fibroblasts

Mouse embryonic fibroblasts (MEFs) were isolated from E13.5 embryos according to standard protocols. Briefly, timed-pregnant females were euthanized, and embryos were harvested under sterile conditions. Embryonic heads and internal organs were removed, and the remaining tissue was minced finely with sterile scissors. Tissue fragments were incubated in 0.25% trypsin-EDTA at 37°C for 15 minutes with periodic agitation. The cell suspension was then neutralized with DMEM containing 10% FBS, passed through a 70 μm cell strainer to remove debris, and centrifuged at 300 × g for 5 minutes. The cell pellet was resuspended in MEF culture medium consisting of DMEM with L-glutamine (Gibco) supplemented with 10% FBS, 1× non-essential amino acids, and 1× penicillin-streptomycin. Primary MEFs were plated in tissue culture flasks and cultured at 37°C in 5% CO_2_. Primary MEFs were immortalized using the NIH 3T3 protocol, passaging cells every 3 days at a constant density of 3 × 10⁵ cells per 60 mm dish for 20-30 passages. For clonal selection, immortalized MEFs were seeded by limiting dilution at 0.5 cells per well in 96-well plates. Wells containing single cells were monitored and expanded over 2-3 weeks.

### Colony Formation Assay

Cells were trypsinized, counted using a hemocytometer, and seeded at low density (1000 cells/plate) in 10 cm plates in standard MEF culture medium. Cells were cultured for 10-14 days at 37°C in 5% CO_2_ with medium changes every 3 days to allow colony formation. After the incubation period, medium was aspirated, and cells were washed twice with phosphate-buffered saline (PBS). Colonies were fixed for 15 minutes at room temperature, then washed twice with PBS. Fixed colonies were stained with 0.5% crystal violet solution for 20 minutes at room temperature. Excess stain was removed by washing plates extensively with distilled water, and plates were air-dried overnight. For each experimental condition, at 3 technical replicates were analyzed, and experiments were repeated in triplicate. Colony formation units were counted per plate. Statistical comparisons between genotypes were performed using unpaired two-tailed Student’s t-tests, with p < 0.05 considered statistically significant.

## Author Contributions

E.P., D.L., E.M.P., and V.L. conceived and designed the project. E.P., D.B., Y.J., and S.Y. performed experiments. E.P. and E.M.P. conducted bioinformatic analyses. E.P., E.M.P., D.L., Y.J., A.S., and V.L. contributed to data interpretation. E.P. and E.M.P. wrote the manuscript with input from all authors. All authors reviewed the manuscript. V.L. provided overall supervision.

## Acknowledgements

We thank the veterinary and technical staff at 14DNR (NIAID) for animal care; the Division of Veterinary Resources (DVR) for pathology services; the Mouse Genetics and Gene Modification (MGGM, NIAID) section for reproductive phenotyping; and the Research Technologies Branch (RTB, NIAID) and the Center for Cancer Research Sequencing Facility (CCR-SF, NCI) for sequencing and data analysis support. This research was supported by the Intramural Research Program of the National Institutes of Health (NIH). The contributions of the NIH author(s) were made as part of their official duties as NIH federal employees, are incompliance with agency policy requirements, and are considered Works of the United States Government. However, the findings and conclusions presented in this paper are those of the author(s) and do not necessarily reflect the views of the NIH or the U.S. Department of Health and Human Services.

## Data Availability

All next-generation sequencing data have been deposited in the Gene Expression Omnibus (GEO) under accession number GSE291162

