## Supplementary Data for "Incomplete developmental silencing of cancer-testis antigen BORIS in humanized mouse model promotes cancer susceptibility"

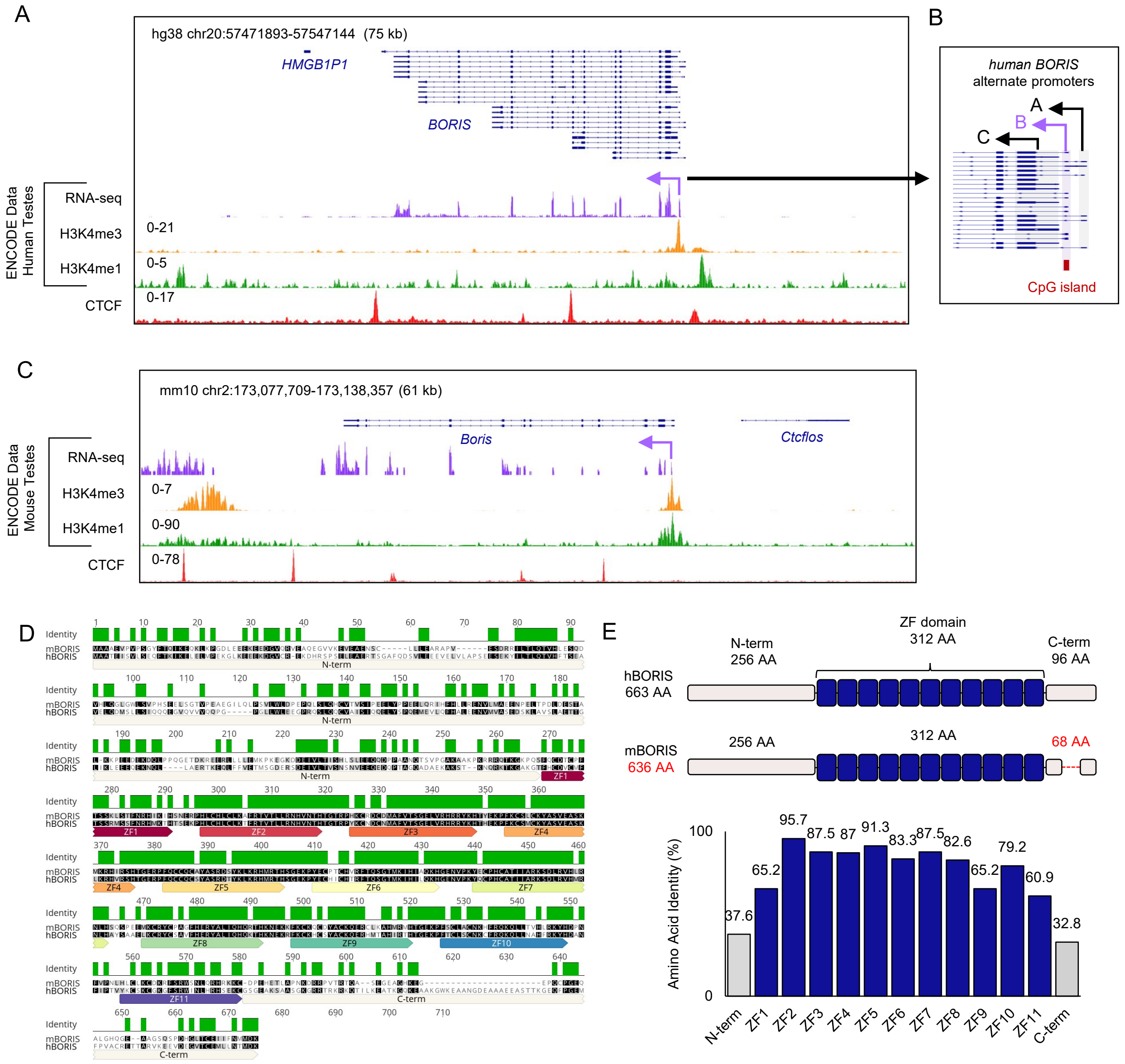


Figure S1. Comparative genomic and protein architecture of human BORIS and mouse Boris.

Figure S1. Comparative genomic and protein architecture of human BORIS and mouse Boris.

(A) Genome browser view of the human *BORIS* locus showing the full 75-kb human sequence inserted into the mouse genome (see Fig. 1). The RefSeq track (dark blue) depicts all 22 human *BORIS* transcript isoforms. The downstream pseudogene *HMGB1P1* is also included in this interval. ENCODE human testes tracks display RNA-seq (purple), H3K4me3 (yellow), H3K4me1 (green), and CTCF ChIP-seq (red); numbers on tracks indicate signal intensity.

(B) Promoter-level view highlighting the three human *BORIS* promoters (A, B, C) and the CpG island uniquely present at promoter B. This regulatory feature is absent from the mouse *Boris* locus.

(C) Genome browser view of the endogenous mouse *Boris* locus, corresponding to the 61-kb region excised in the humanization strategy (see Fig. 1). The RefSeq track shows the two annotated mouse transcripts arising from a single promoter. ENCODE mouse testes datasets show RNA-seq (purple), H3K4me3 (yellow), H3K4me1 (green), and CTCF ChIP-seq (red); numbers indicate signal intensity.

(D) Protein sequence alignment of hBORIS and mBORIS generated using Geneious Prime, using canonical amino acid sequences from Ensembl. N-terminal region, zinc-fingers 1–11, and C-terminal domains are annotated below the aligned sequences.

(E) Schematic representation of human and mouse BORIS protein structure, comparing relative lengths of the N-terminal, ZF array, and C-terminal domains. The accompanying bar chart shows average amino acid identity (%) for each domain, as quantified from the alignment in panel D.

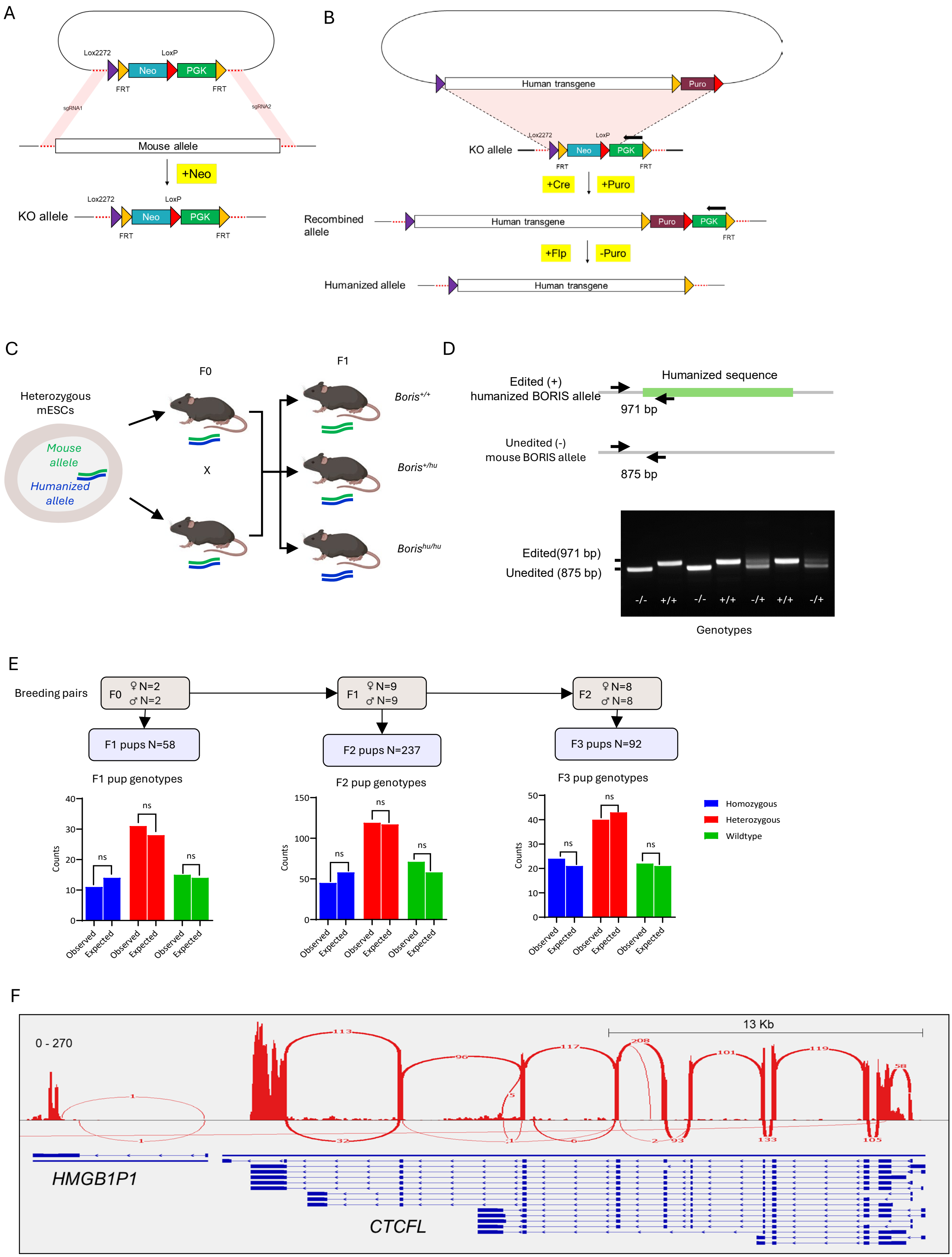


Figure S2. Generation and validation of the humanized BORIS mouse line

Figure S2. Generation and validation of the humanized BORIS mouse line

(A) CRISPR-mediated deletion of the endogenous mouse Boris locus. Two CRISPR/Cas9 guide RNAs were designed to excise the 61-kb mouse Boris genomic interval (see Fig. 1 and Fig. S1). Homology-directed repair was driven using a short donor construct containing a PGK-Neo selection cassette, resulting in precise replacement of the endogenous locus with Neo, thereby generating a "landing pad" allele for recombinase-mediated genomic replacement (RMGR). Step 2 of RMGR showing construct containing 75 kb human sequence BORIS locus engineered from donor human BAC Cre inserted into target site in knocked out sequence. Puro and flip are used to remove target sequences.

(B) RMGR with the full 75-kb human BORIS locus. A BAC-derived RMGR donor construct containing the entire 75-kb human BORIS genomic region—spanning all promoters (A, B, and C), exons, introns, and regulatory elements—was targeted into the Neo-disrupted locus by CRISPR-assisted HDR. The donor cassette carried Puro and PGK selection modules flanked by loxP and FRT sites. Sequential Cre- and Flp-mediated excision removed both selection cassettes, generating a seamless humanized allele containing only the human genomic sequence flanked by native mouse chromosomal context.

(C) Derivation of heterozygous humanized mESC clones and establishment of mouse lines. Correctly targeted mESCs carrying one humanized allele were used to generate F1 founder mice, which were intercrossed to establish homozygous humanized, heterozygous, and wild-type mice. The schematic illustrates allele transmission across generations.

(D) Genotyping strategy and representative PCR validation. Alleles were distinguished by amplicon size: the 971-bp product corresponds to the humanized allele, and the 875-bp product to the unaltered mouse allele. Shown are representative agarose gels illustrating clear discrimination of wild-type ($Boris^{+/+}$), heterozygous ($Boris^{+/hu}$), and homozygous humanized ($Boris^{hu/hu}$) mice.

(E) Mendelian transmission across F0–F3 generations. Breeding outcomes from F0, F1, and F2 heterozygous intercrosses are summarized. All pups were genotyped, and observed allele frequencies (absolute counts) were compared with expected Mendelian ratios using chi-square testing. Bar plots show homozygous humanized (blue), heterozygous (red), and wild-type (green) distributions. NS denotes no significant deviation from Mendelian expectations (p-value <0.05).

(F) RNA-seq confirmation of human promoter usage in humanized BORIS mouse testes. RNA-seq coverage tracks from $\text{Boris}^{hu/hu}$ testes demonstrate robust transcription from canonical human promoter B (see Fig. S1B), confirming correct promoter deployment and physiological transcriptional activity of the humanized locus *in vivo*.

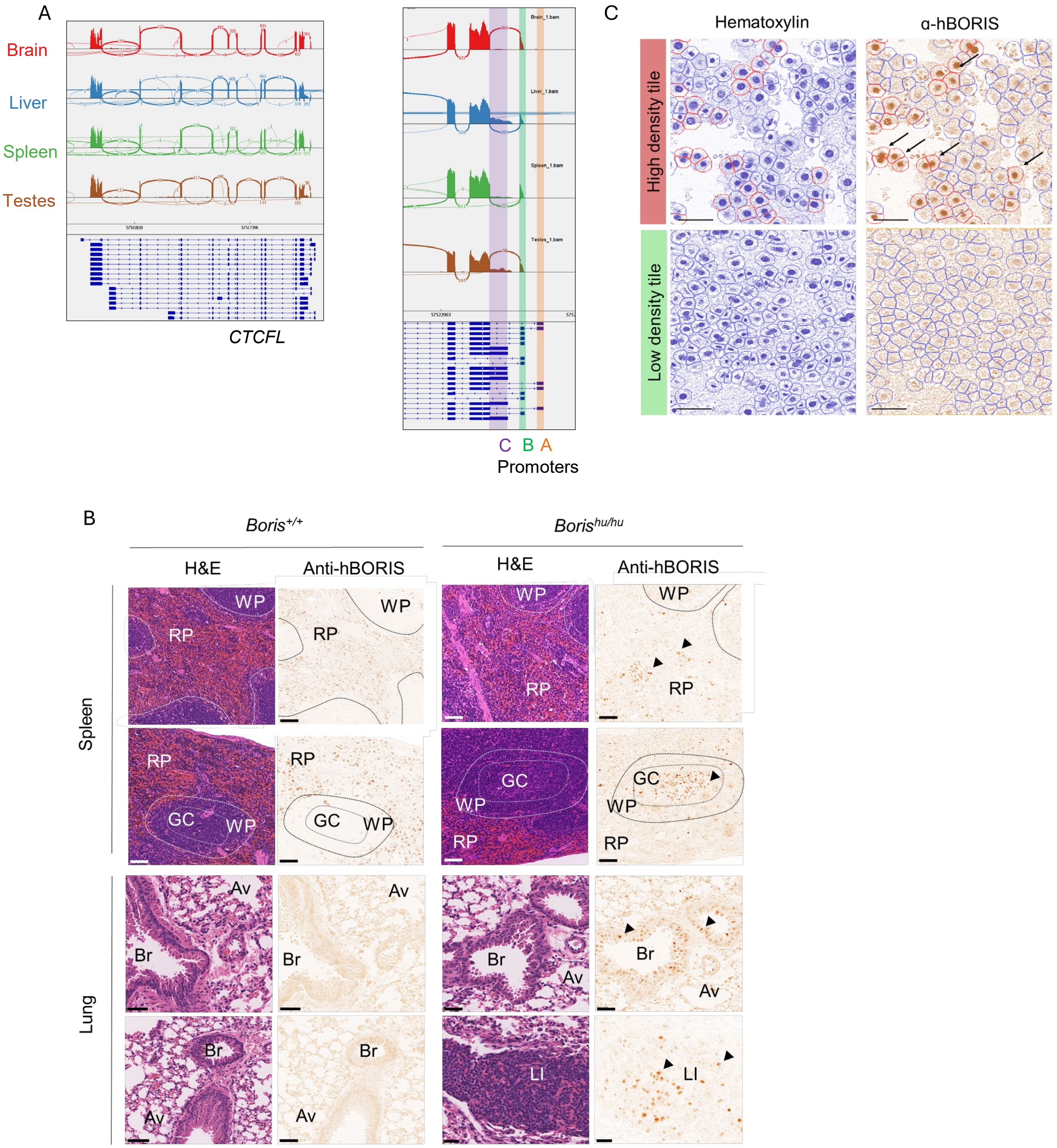


Figure S3. Mosaic hBORIS expression in somatic tissues with alternative promoter usage.

Figure S3. Mosaic hBORIS expression in somatic tissues with alternative promoter usage.

(A) RNA-seq coverage tracks across the human *BORIS* locus in *Boris*$^{hu/hu}$ somatic tissues. Somatic tissues predominantly utilize promoter B (purple box), with minor promoter C activity in liver (green box). Promoter A (orange box) remains silent. Heatmap shows read density across transcript isoforms.

(B) Anti-hBORIS immunohistochemistry in *Boris*$^{hu/hu}$ spleen and lung reveals heterogeneous expression (arrowheads). BORIS-positive cells localize to red pulp (RP), white pulp (WP), and germinal centers (GC) in spleen; sparse staining appears in bronchi (Br), alveoli (Av), and lymphoid infiltrates (LI) in lung. *Boris*$^{+/+}$ controls show no signal. Scale bars, 50 μm.

(C) QuPath automated tiling quantification method. Tissue divided into equal tiles scored for BORIS-positive nuclei (brown DAB). High-density tiles (red bar) show clustered expression; low-density tiles (green bar) show sparse staining. Hematoxylin counterstain (blue). Scale bar, 50 μm. Quantification in Fig 2C.

A

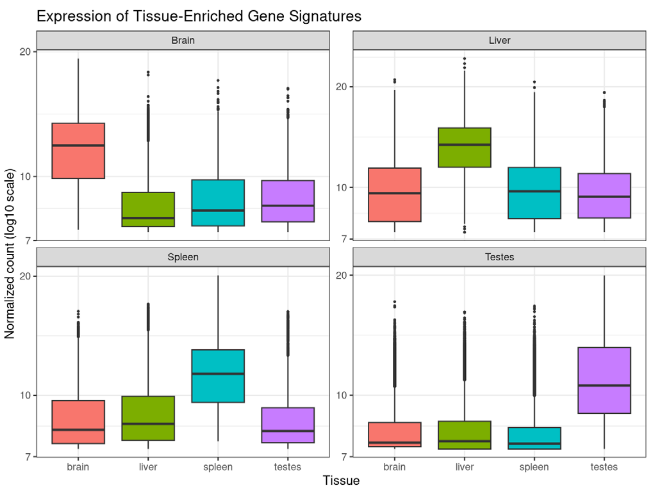


B

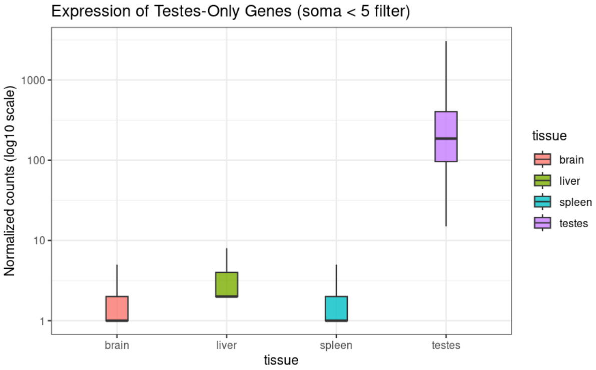


| Number of tissue-enriched signature genes (WT samples) | |
|---|---|
| Tissue | N_Signature_Genes |
| Brain | 3380 |
| Liver | 2495 |
| Spleen | 2256 |
| Testes (TEGs) | 7053 |

C

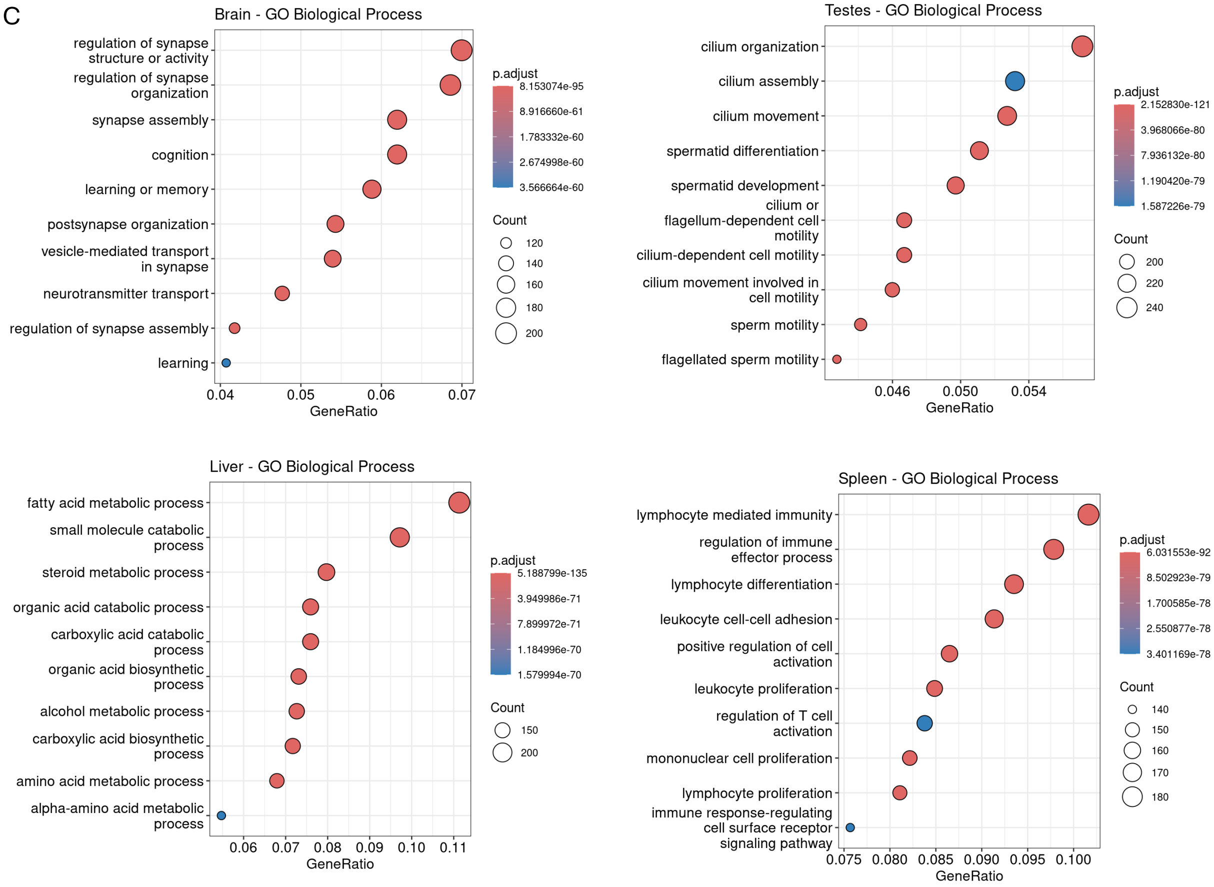


Figure S4. Tissue-enriched gene signatures in wildtype mice.

Figure S4. Tissue-enriched gene signatures in wildtype mice.

(A) Expression distribution of tissue-enriched genes across wildtype tissues. Box plots show each signature's expression pattern: brain (n=3,380 genes), liver (n=2,495), spleen (n=2,256), and testes (n=7,053 TEGs). Testes-enriched genes show marked elevation in testes relative to somatic tissues. RNA-seq from *Boris*$^{+/+}$ mice, 16 weeks, n=2 per tissue.

(B) Testes-specific genes (TSGs, n=2,215, defined by soma <5 normalized counts) show >100-fold enrichment in testes. Table summarizes tissue-enriched signature sizes used as reference frameworks for identifying ectopic activation in *Boris*$^{hu/hu}$ tissues.

(C) GO Biological Process enrichment confirms functional specificity of tissue signatures. Brain: neuronal processes (synapse regulation, cognition, learning). Testes: spermatogenesis (cilium assembly, spermatid differentiation, sperm motility). Liver: metabolism (fatty acid, steroid pathways). Spleen: immunity (lymphocyte activation, T cell function). Dot size = gene count; color = adjusted p-value. Only terms with FDR <0.05 shown.

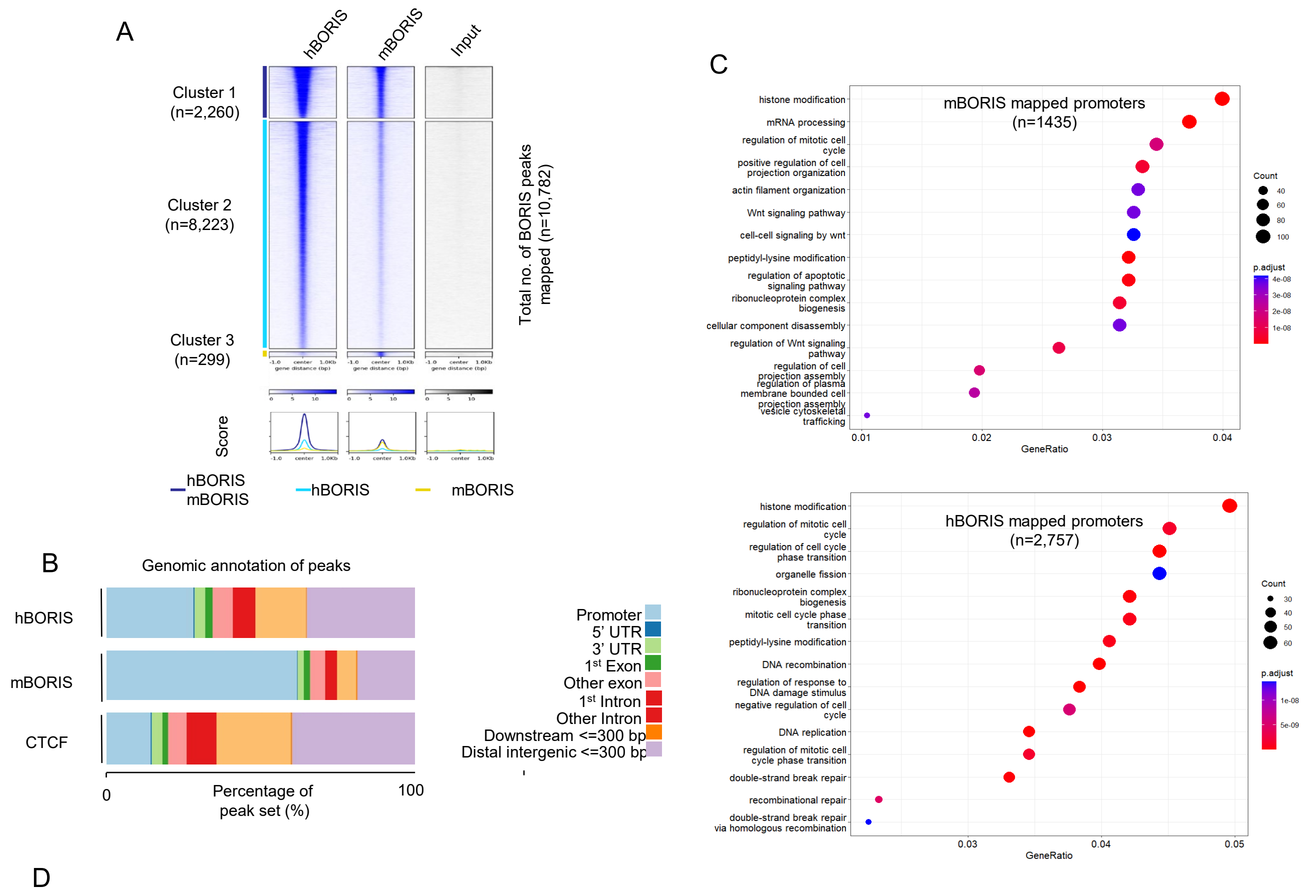


D

| Ab | Rep | Reads | Dup% | Read length | Frag. length | RelCC | SSD | RiP% |
|---|---|---|---|---|---|---|---|---|
| hBORIS | 1 | 1843929 | 0 | 35 | 159 | 6.4 | 1.3 | 2.8 |
| hBORIS | 2 | 2282547 | 0 | 35 | 161 | 9.2 | 1 | 1.1 |
| hBORIS | 3 | 1782086 | 0 | 35 | 166 | 7.1 | 1 | 1.5 |
| mBORIS | 1 | 831056 | 0 | 48 | 159 | 0.62 | 1.1 | 0.61 |
| mBORIS | 2 | 1219784 | 0 | 48 | 163 | 0.78 | 1.1 | 0.66 |
| NA | 1 | 1495365 | 0 | 35 | 127 | 10 | 0.82 | 0.16 |
| NA | 1 | 811076 | 0 | 48 | 171 | 2.4 | 1.3 | 0.095 |

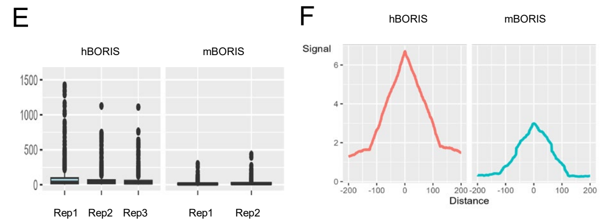


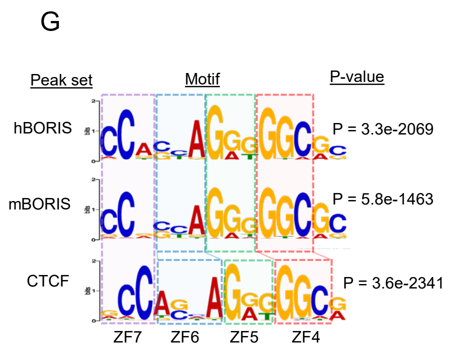


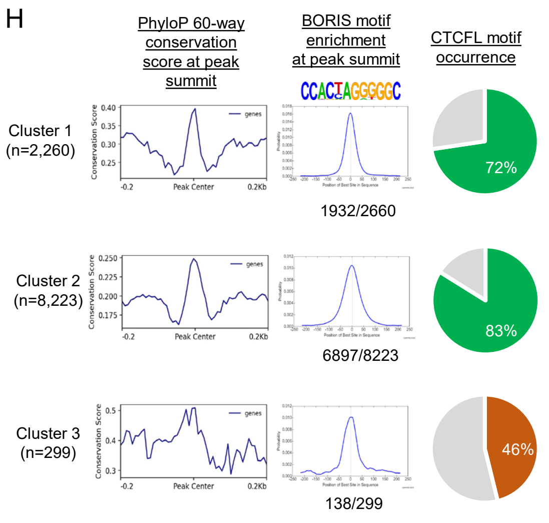


Figure S5. ChIP-seq quality metrics and binding site characterization in testes.

Figure S5. ChIP-seq quality metrics and binding site characterization in testes.

(A) Comparison of hBORIS ChIP-seq in humanized testes with published mBORIS datasets. Heatmaps show signal ±1 kb from peak summits for three clusters: Cluster 1 (n=2,260) = conserved binding sites; Cluster 2 (n=8,223) = expanded detection in hBORIS; Cluster 3 (n=299) = weak sites. Aggregate profiles below.

(B) Genomic distribution of hBORIS peaks. Enrichment at promoters (light blue), UTRs (dark blue/purple), and exons (green/yellow). mBORIS and CTCF show similar distributions.

(C) GO enrichment of genes with promoter-proximal BORIS binding. mBORIS (n=1,435 genes): chromatin organization, histone modification. hBORIS (n=2,757 genes): chromatin processes plus germline-specific terms (cilium organization, meiosis, DNA recombination, sperm development). Dot size = gene count; color = adjusted p-value.

(D) Fragment length distribution confirms enrichment (~150-200 bp) in hBORIS and mBORIS ChIP-seq across replicates.

(E) Cross-coverage plots show read density ±500 bp from peak centers. Both hBORIS (red) and mBORIS (cyan) show central enrichment, confirming specific binding.

(F) ChIPQC quality metrics. hBORIS: 1.8-2.3M reads, RSC 6.4-7.1, RiP% 1.1-2.8, exceeding ENCODE standards. mBORIS: lower depth and RSC values.

(G) De novo motif discovery identifies canonical CTCF binding motif at peak summits for hBORIS, mBORIS, and CTCF. Near-identical PWMs confirm conserved DNA recognition via zinc finger domain (ZF7-ZF4 modules shown).

(H) Evolutionary conservation and motif occurrence at BORIS binding sites. PhyloP 60-way conservation scores (left) show elevated conservation at peak summits. BORIS motif enrichment (middle) and percentage of peaks containing canonical CTCF motif (right): Cluster 1: 72% (1,932/2,660); Cluster 2: 83% (6,897/8,223); Cluster 3: 46% (138/299). High motif occurrence validates authentic target site recognition.

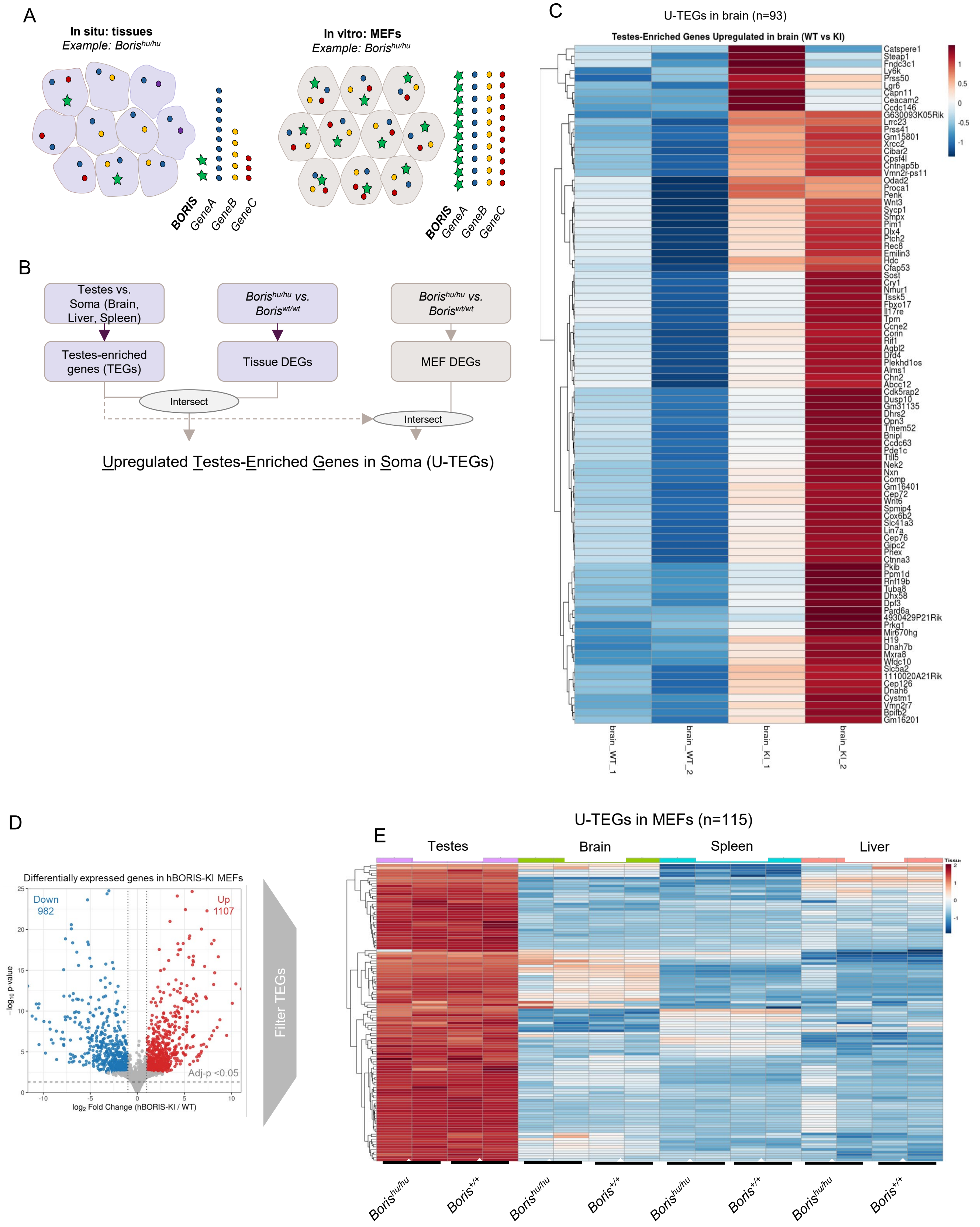


Figure S6. Identification of upregulated testes-enriched genes (U-TEGs) in BORIS-expressing somatic contexts.

Figure S6. Identification of upregulated testes-enriched genes (U-TEGs) in BORIS-expressing somatic contexts.

(A) Schematic illustrating analytical challenge of mosaic BORIS expression. Left (in situ tissues): Heterogeneous tissues contain mixed populations with few BORIS-positive cells (green stars) among BORIS-negative cells (colored dots), diluting transcriptional signatures in bulk RNA-seq. Right (in vitro MEFs): Clonally selected BORIS-positive MEF lines may provide more homogeneous populations for clearer signal detection, though BORIS expression levels vary between clones. Combined analysis of both contexts enables robust U-TEG identification despite cellular heterogeneity.

(B) Bioinformatic pipeline for U-TEG identification. Testes-enriched genes (TEGs, n=7,053) defined by comparing wildtype testes vs somatic tissues (brain, liver, spleen). Differentially expressed genes (DEGs) identified in $\textit{Boris}^{hu/hu}$ vs $\textit{Boris}^{+/+}$ tissues and MEFs. Intersection of upregulated DEGs with TEG reference signature defines U-TEGs: genes normally enriched in testes that become ectopically activated in BORIS-expressing somatic contexts.

(C) Heatmap of U-TEGs in $\textit{Boris}^{hu/hu}$ brain (n=93 genes). Expression shown across genotypes and tissues. U-TEGs exhibit low/absent expression in wildtype brain but show upregulation in $\textit{Boris}^{hu/hu}$ brain, with coordinate high expression in testes (red). Hierarchical clustering groups genes by expression pattern. Gene names listed at right; selected examples include meiotic and germline-restricted genes.

(D) Volcano plot of differentially expressed genes in BORIS-positive vs BORIS-negative MEF clones. Upregulated genes (red, n=1,107) include U-TEGs; downregulated genes (blue, n=982). Dashed lines: adjusted $p<0.05$ and $|\log_2 FC|>1$ thresholds.

(E) Heatmap of U-TEGs identified in MEFs (n=115 genes) showing expression across testes, brain, spleen, and liver in both $\textit{Boris}^{hu/hu}$ and $\textit{Boris}^{+/+}$ mice. U-TEGs show high testes expression (red) and ectopic activation in $\textit{Boris}^{hu/hu}$ MEFs, brain, spleen, and/or liver (variable patterns), while remaining low in $\textit{Boris}^{+/+}$ somatic tissues (blue). "Filter TEGs" label indicates these represent the subset of TEGs showing upregulation in BORIS-expressing contexts. Expression normalized within genes (row z-scores); hierarchical clustering groups genes by pattern.

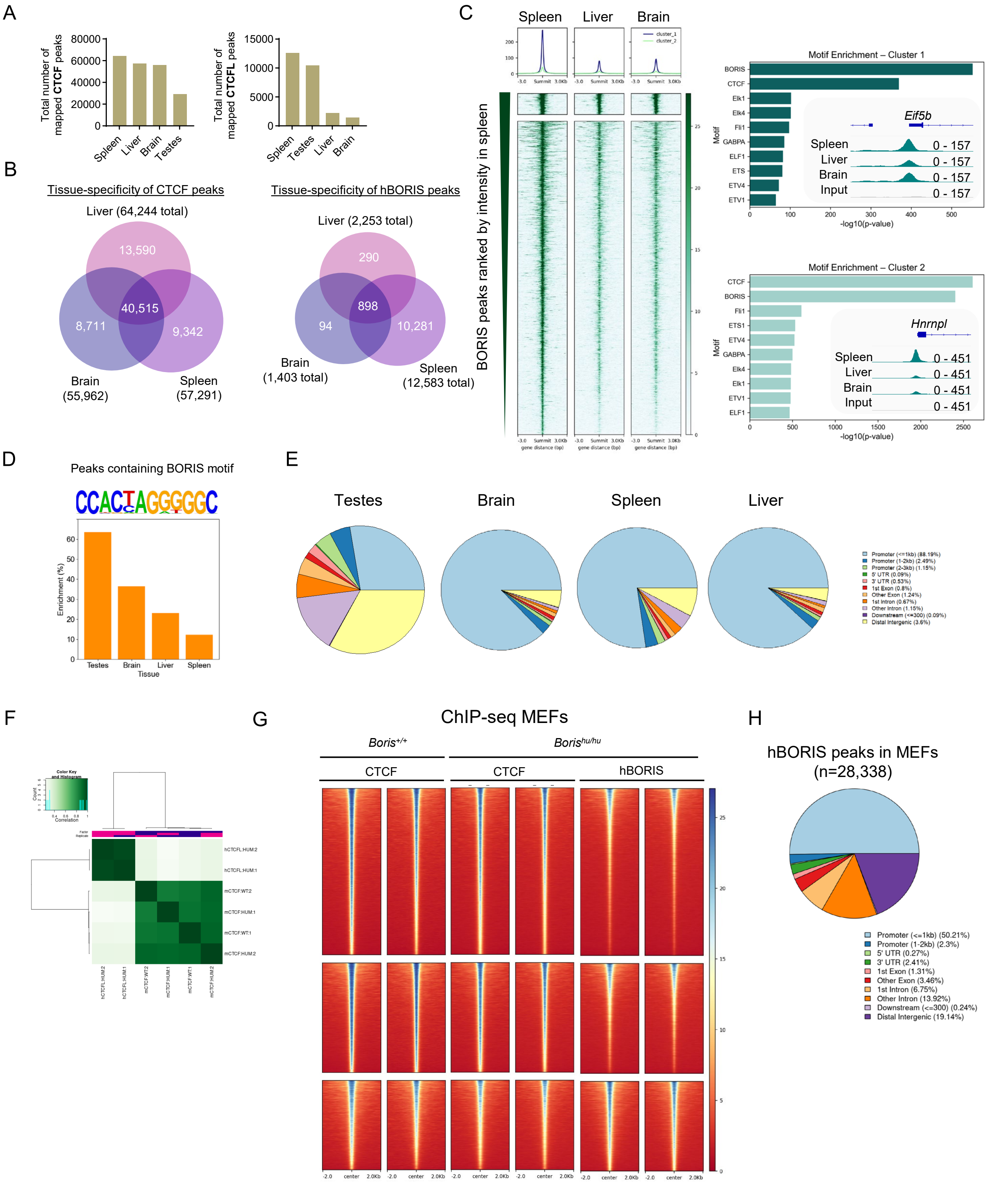


Figure S7. ChIP-seq characterization of hBORIS and CTCF binding in somatic tissues and MEFs.

Figure S7. ChIP-seq characterization of hBORIS and CTCF binding in somatic tissues and MEFs.

(A) Total hBORIS (n=12,583 spleen, 2,252 liver, 1,403 brain) and CTCF peaks across *Boris*$^{hu/hu}$ tissues.

(B) Venn diagrams show tissue-specific vs shared hBORIS (left) and CTCF (right) peaks. CTCF shows extensive overlap (40,515 shared sites), while hBORIS exhibits greater tissue specificity.

(C) Heatmaps and genome browser tracks show tissue-enriched hBORIS binding in spleen, liver, and brain (±1.5 kb from summits). Density profiles above heatmaps. Motif enrichment analysis (right panels) for spleen-enriched clusters identifies canonical CTCF/BORIS motif and tissue-specific co-factors (e.g., *Etf5b* for Cluster 1, *Hnrnpl* for Cluster 2).

(D) BORIS motif logo and enrichment frequency across tissues; highest in testes (>60%), lower in soma (~20-35%).

(E) Genomic distribution pie charts. Testes shows diverse distribution (promoters, introns, intergenic). Somatic tissues show increased promoter-proximal binding (light blue), particularly in brain, spleen, and liver (>50% promoter/5'UTR/3'UTR).

(F) Pearson correlation heatmap demonstrates high reproducibility between hBORIS ChIP-seq biological replicates in MEFs (r>0.9).

(G) Signal heatmaps (±2 kb from summits) comparing CTCF binding in *Boris*$^{+/+}$ MEFs (left), CTCF in *Boris*$^{hu/hu}$ MEFs (middle), and hBORIS in *Boris*$^{hu/hu}$ MEFs (right). Three biological replicates shown per condition. CTCF signal is maintained in humanized MEFs, with hBORIS showing co-occupancy at CTCF-bound sites.

(H) Genomic distribution of hBORIS peaks in MEFs (n=28,338). Promoter-enriched (light blue, 29.5%), with substantial 5'UTR (dark blue, 2.3%), intergenic (purple, 24%), and intronic binding (orange, 13.92%; brown, 3.24%). Distribution differs from somatic tissues, reflecting homogeneous BORIS-positive cell population.

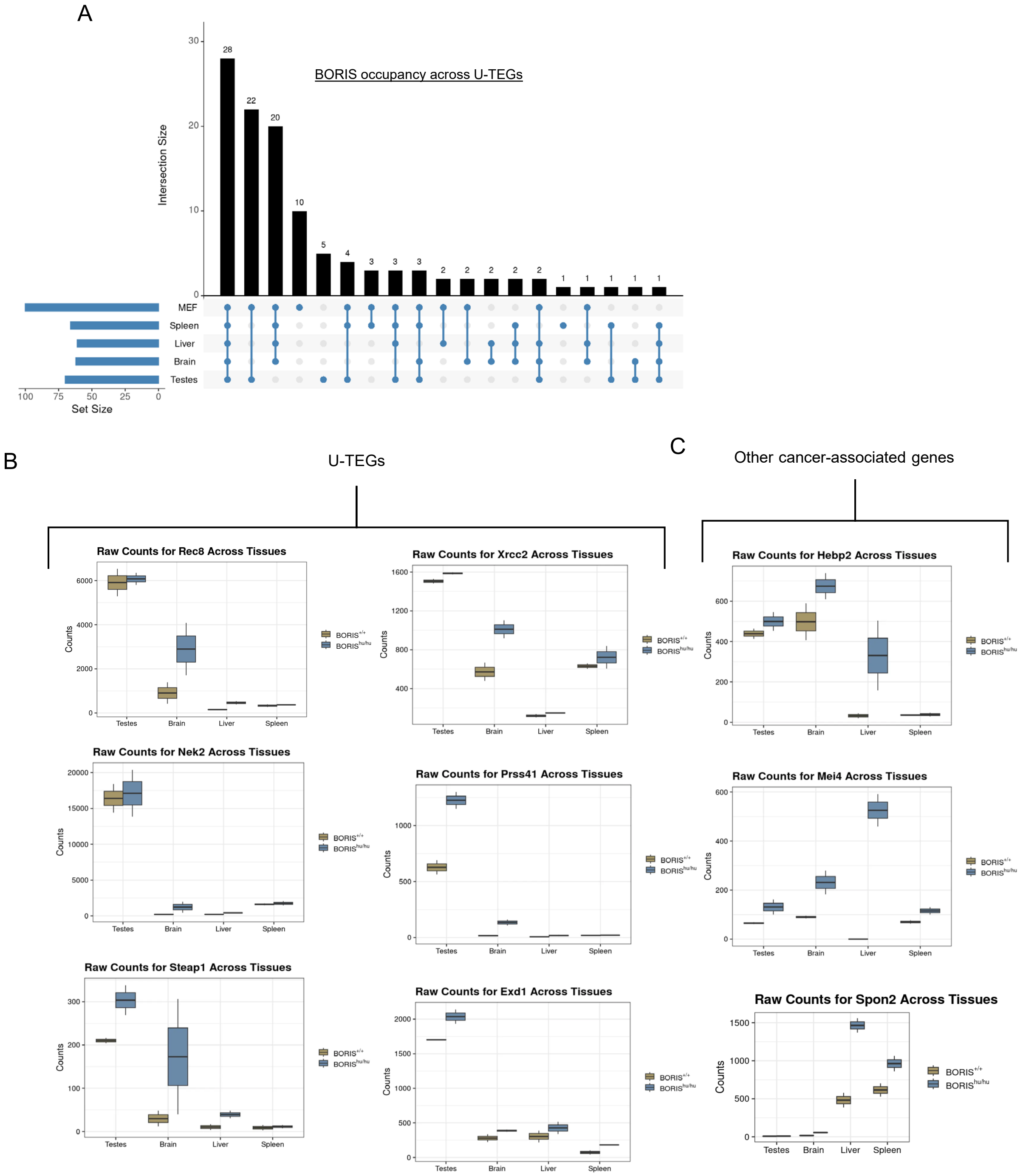


Figure S8. BORIS occupancy at U-TEG promoters and expression of cancer-associated genes.

Figure S8. BORIS occupancy at U-TEG promoters and expression of cancer-associated genes.

(A) UpSet plot showing BORIS binding across U-TEGs in different somatic contexts. Top: Bar graph displays intersection size (number of U-TEGs with BORIS occupancy across tissue combinations). Largest intersections: 28 U-TEGs bound in MEFs only, 22 in MEFs+spleen, 20 in MEFs+spleen+liver. Bottom: Horizontal bars show total U-TEG counts per tissue: MEFs (n=115), spleen (n=13), liver (n=12), brain (n=93), testes (germline reference). Connected dots indicate tissue combinations for each intersection. Overall, 108 of 197 analyzable U-TEGs (55%) show detectable BORIS binding in at least one somatic context, with MEFs showing highest detection due to homogeneous BORIS expression.

(B) Representative U-TEGs showing tissue-specific upregulation in $Boris^{hu/hu}$ mice. Box plots display RNA-seq raw counts across testes, brain, liver, and spleen for $Boris^{+/+}$ (tan) and $Boris^{hu/hu}$ (blue). Rec8: Meiotic cohesin component, high in testes, upregulated in $Boris^{hu/hu}$ brain and liver. Xrcc2: DNA recombination factor, testes-enriched, activated in $Boris^{hu/hu}$ brain and spleen. Nek2: Kinase involved in meiosis, shows testes-specific expression with modest brain activation. Prss41: Serine protease, testes-restricted, upregulated in $Boris^{hu/hu}$ brain. Steap1: Metalloreductase, elevated in $Boris^{hu/hu}$ brain. Exd1: Exonuclease, activated across multiple $Boris^{hu/hu}$ tissues. All show baseline low/absent expression in $Boris^{+/+}$ soma (n=2 biological replicates per genotype).

(C) Cancer-associated genes upregulated in $Boris^{hu/hu}$ somatic tissues. Hebp2: Heme-binding protein implicated in cancer pathways, upregulated in $Boris^{hu/hu}$ liver and spleen. Mei4: Elevated in $Boris^{hu/hu}$ spleen. Spon2: Extracellular matrix protein, activated in $Boris^{hu/hu}$ liver and spleen. While not uniformly testes-enriched, these genes show ectopic activation in BORIS-expressing contexts, suggesting broader transcriptional dysregulation beyond U-TEGs (n=2 per genotype).

A

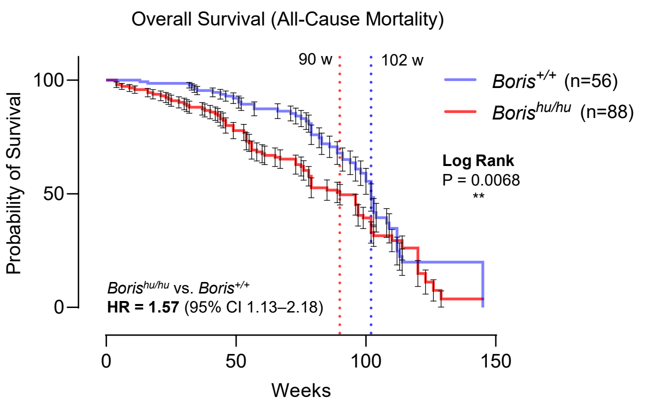


B

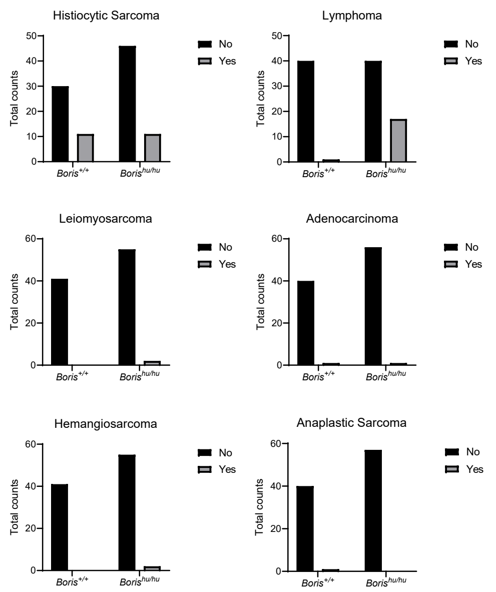


C

Contingency Tables Used for Fisher's Exact Tests

| Tumor Type | Genotype | No | Yes | Total |
|---|---|---|---|---|
| **Histiocytic Sarcoma** | *Boris*$^{+/+}$ | 30 | 11 | 41 |
| | *Boris*$^{hu/hu}$ | 46 | 11 | 57 |
| **Lymphoma** | *Boris*$^{+/+}$ | 40 | 1 | 41 |
| | *Boris*$^{hu/hu}$ | 40 | 17 | 57 |
| **Leiomyosarcoma** | *Boris*$^{+/+}$ | 41 | 0 | 41 |
| | *Boris*$^{hu/hu}$ | 55 | 2 | 57 |
| **Adenocarcinoma** | *Boris*$^{+/+}$ | 40 | 1 | 41 |
| | *Boris*$^{hu/hu}$ | 57 | 0 | 57 |
| **Hemangiosarcoma** | *Boris*$^{+/+}$ | 41 | 0 | 41 |
| | *Boris*$^{hu/hu}$ | 55 | 2 | 57 |
| **Anaplastic Sarcoma** | *Boris*$^{+/+}$ | 40 | 1 | 41 |
| | *Boris*$^{hu/hu}$ | 57 | 0 | 57 |

D

Summary of Tumor Incidence by Genotype

| Tumor Type | *Boris*$^{+/+}$ | *Boris*$^{hu/hu}$ | Fisher's Exact P-value | Significance |
|---|---|---|---|---|
| **Histiocytic Sarcoma** | 11/41 (26.8%) | 11/57 (19.3%) | 0.4636 | ns |
| **Lymphoma** | 1/41 (2.4%) | 17/57 (29.8%) | **0.0004** | *** |
| **Leiomyosarcoma** | 0/41 (0%) | 2/57 (3.5%) | 0.5083 | ns |
| **Adenocarcinoma** | 1/41 (2.4%) | 0/57 (0%) | 0.4184 | ns |
| **Hemangiosarcoma** | 0/41 (0%) | 2/57 (3.5%) | 0.5083 | ns |
| **Anaplastic Sarcoma** | 1/41 (2.4%) | 0/57 (0%) | 0.4184 | ns |

E

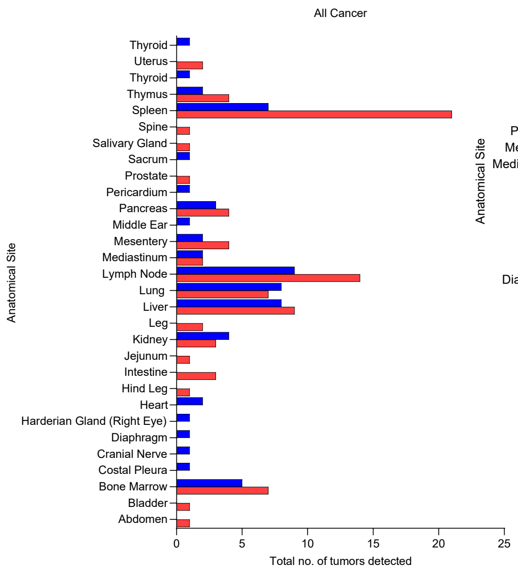


F

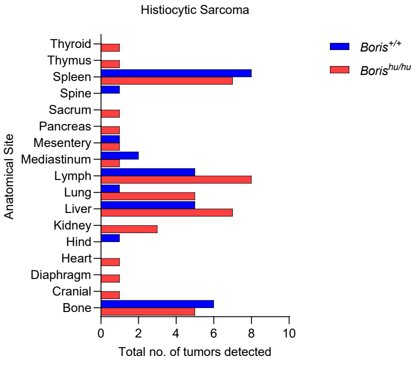


G

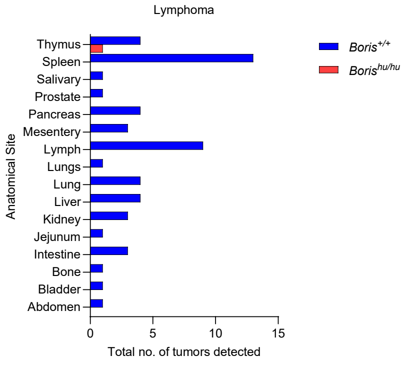


Figure S9. Tumor spectrum, incidence, and anatomical distribution in *Boris*$^{+/+}$ and *Boris*$^{hu/hu}$ mice.

Figure S9. Tumor spectrum, incidence, and anatomical distribution in *Boris*$^{+/+}$ and *Boris*$^{hu/hu}$ mice.

(A) Kaplan–Meier analysis of overall survival for *Boris*$^{+/+}$ (n = 56) and *Boris*$^{hu/hu}$ (n = 88) mice. *Boris*$^{hu/hu}$ animals exhibited reduced survival relative to *Boris*$^{+/+}$, with a hazard ratio of 1.57 (95% CI, 1.13–2.18). Vertical dashed lines indicate median survival for each genotype (90 and 102 weeks). Survival curves were compared by log-rank test (P = 0.0068).

(B) Distribution of necropsy-confirmed tumor types in *Boris*$^{+/+}$ and *Boris*$^{hu/hu}$ mice. Bar plots show the total number of animals within each genotype classified as tumor-positive or tumor-negative for histiocytic sarcoma, lymphoma, leiomyosarcoma, adenocarcinoma, hemangiosarcoma, and anaplastic sarcoma. Counts reflect categorical incidence and were used for subsequent statistical comparisons.

(C) Contingency tables used for Fisher's exact tests comparing tumor incidence between *Boris*$^{+/+}$ and *Boris*$^{hu/hu}$ cohorts. Tables report the number of animals with or without each tumor type and total animals evaluated per genotype for histiocytic sarcoma, lymphoma, leiomyosarcoma, adenocarcinoma, hemangiosarcoma, and anaplastic sarcoma. These values were used to generate statistical results shown panel C.

(D) Summary of tumor incidence by genotype. Table reports the proportion of *Boris*$^{+/+}$ (n = 41) and *Boris*$^{hu/hu}$ (n = 57) animals developing each tumor type, along with corresponding Fisher's exact test P-values. Lymphoma incidence was significantly elevated in *Boris*$^{hu/hu}$ mice (P = 0.0004), whereas no significant genotype-dependent differences were observed for other tumor types (ns).

(E) (E-G) Anatomical distribution of all tumors (D), histiocytic sarcomas (E) and lymphomas (F) detected in *Boris*$^{+/+}$ and *Boris*$^{hu/hu}$ mice. Bar plots show the total number of lesions identified at each site, illustrating increased tumor burden and broader tissue involvement in *Boris*$^{hu/hu}$ animals.

Table S1. PacBio long-read validation of the humanized BORIS transgene.

Summary of PacBio HiFi long-read sequencing confirming the structural integrity of the 75-kb human BORIS locus inserted into the mouse genome. The table reports all detected sequence features across the transgene, including genomic position, allele, variant consequence, HGVS annotations, predicted functional impact, population allele frequencies, SpliceAI and CADD scores, and relevant regulatory or phenotypic annotations.

Table S2. Tissue-Enriched Gene Signatures, Differentially Expressed Genes, and Upregulated Testes-Enriched Genes.

This table compiles RNA-seq–derived gene sets used to characterize tissue specificity and BORIS-dependent transcriptional changes.

Table S3. Tumor Pathology Data

Complete pathology data for survival cohort monitored until 2 years age or endpoint morbidity.
